# Cf-4- and Cf-5-Triggered Plant Immunity: Similarities and Differences

**DOI:** 10.64898/2026.06.10.731257

**Authors:** Esranur Budak, Henrique Aguiar Canha, Matthieu H.A.J. Joosten

## Abstract

Plant immunity is, amongst others, mediated by receptor like proteins (RLPs), which are localized on the plasma membrane. RLPs recognize extracellular immunogenic patterns (ExIPs) originating from pathogens or derived from the host itself, which leads to extracellularly triggered immunity (ExTI). Cf proteins, which are well-known RLPs of tomato (*Solanum lycopersicum*) confer resistance against the fungal pathogen *Fulvia fulva.* Cf-9, Cf-4, Cf-2 and Cf-5 are well-known examples of Cf proteins, mediating recognition of the matching ExIPs Avr9, Avr4, Avr2 and Avr5, respectively, which are secreted effectors of *F. fulva* and trigger hypersensitive response (HR)-related cell death in tomato plants carrying these Cf proteins. Although all these Cf proteins confer proper resistance to the fungus, Cf-9 and Cf-4 trigger a stronger and faster cell death than Cf-5 and Cf-2. It is unknown whether these phenotypical differences arise from variations in the molecular mechanism of the cellular immune response that is initiated by the Cf proteins, and whether this phenotypic difference correlates with varying degrees in the intensity and timing of the triggered immune responses and robustness of the resistance. To try to answer these questions, in this study the immune responses triggered by Cf-4 and Cf-5 were compared. Cf-4 and Cf-5 share the same core upstream signaling components to trigger HR-related cell death in *Nicotiana benthamiana*. In tomato, both receptors induce rapid MAPK activation, which is more sustained for the Cf-5/Avr5 combination. Both Avr4 and Avr5 induce an apoplastic burst of reactive oxygen species (ROS), independently of the presence of their matching receptors, while remaining dependent on RBOHB for this ROS burst. Full transcriptome analysis at 3 and 7 hours after immune activation revealed a large shared set of differentially expressed genes, alongside qualitative and quantitative differences, with the Cf-5/Avr5 combination inducing a broader transcriptional reprogramming. Despite these differences, Cf-4 and Cf-5 confer a comparable level of resistance to *F. fulva.* These results demonstrate that Cf-4 and Cf-5 share conserved immune initiation mechanisms, but diverge in downstream signaling dynamics, and that the intensity and timing of the HR-related cell death do not affect the robustness of the resistance.

## Introduction

Plants are constantly challenged by a wide range of microbial pathogens, and have evolved immune systems to protect themselves against infections. The gene-for-gene model is a well-known concept in plant disease resistance, in which host resistance (*R*) genes mediate recognition of matching avirulence (Avr) genes of the invading pathogen, eventually resulting in the activation of defense responses and disease resistance (Flor, 1971). The interaction between *Fulvia fulva* (syn. *Cladosporium fulvum*) and tomato is a well-known example of a pathosystem following the gene-for-gene model (Joosten and de Wit, 1999; de Wit et al., 2009; de Wit, 2016). *F. fulva* is a non-obligate hemibiotrophic fungal pathogen that causes the tomato leaf mold disease, and resistance to this fungus is mediated by *Cf* genes that have been introgressed from wild tomato species into cultivated tomato. *F. fulva* hyphae enter tomato leaves through stomata of susceptible as well as resistant plants. In the absence of *Cf* genes matching a *F. fulva* effector, the fungus colonizes the apoplast and eventually produces excessive amounts of conidiophores, which clog the stomata and lead to leaf wilting. However, when a matching *Cf*/*Avr* combination is present, a swift defense response is mounted that leads to resistance (Rivas and Thomas, 2005; de Wit, 2016; Zhao et al., 2022).

At the molecular level, plants have a two-layered innate immune system. The first layer of immunity is mediated by cell surface receptors that recognize extracellular immunogenic patterns (ExIPs), leading to extracellularly-triggered immunity (ExTI) (van der Burgh and Joosten, 2019). Cell surface immune receptors are classified into two groups: receptor-like kinases (RLKs) and receptor-like proteins (RLPs). Both receptor types contain an ectodomain, which frequently contains leucine-rich repeat (LRR) motifs that are essential for proteinaceous ligand recognition, and a transmembrane domain. RLKs also have an intracellular kinase domain, whereas RLPs lack such a kinase domain (Böhm et al., 2014; Monaghan and Zipfel, 2012; Ranf, 2017; Zipfel, 2014).

FLAGELLIN SENSING 2 (FLS2) is a well-known example of an LRR-RLK that recognizes the bacterial flagellin-derived peptide FLAGELLIN 22 (flg22) (Gómez-Gómez and Boller, 2000). Upon recognition of flg22, the LRR-RLK SOMATIC EMBRYOGENESIS RECEPTOR KINASE 3/BRI1-ASSOCIATED RECEPTOR KINASE 1 (SERK3/BAK1) is recruited by FLS2, and downstream cytoplasmic immune signaling is initiated by the two cytoplasmic kinase domains that activate each other by trans-phosphorylation (Chinchilla et al., 2007). Cf proteins are LRR-RLP-type cell surface receptors that recognize matching small, secreted, cysteine-rich effector proteins released by *F. fulva* when colonizing the apoplast, rendering these effectors avirulence (Avr) proteins. Because Cf proteins lack a cytoplasmic kinase domain, they constitutively interact with the RLK SUPPRESSOR OF BIR1-1/EVERSHED (SOBIR1/EVR) (Liebrand et al., 2013). Upon recognition of the matching Avr protein, the Cf/SOBIR1 complex also recruits BAK1, here leading to trans-phosphorylation events between the cytoplasmic kinase domains of SOBIR1 and BAK1 and subsequent phosphorylation of downstream immune signaling partners (van der Burgh et al., 2019; Huang et al., 2024; Postma et al., 2016). This signaling cascade initiates downstream immune responses, including a burst of apoplastic reactive oxygen species (ROS), the activation of mitogen-activated protein kinases (MAPKs), transcriptional regulation of immune-related genes, and finally the HR associated with programmed cell death (PCD), all together eventually resulting in resistance to *F. fulva* (Huang and Joosten, 2025).

Among the various *Cf* resistance genes that have been identified, *Cf-4*, *Cf-9*, *Cf-2*, and *Cf-5* can be classified into two groups, based on their sequence and their specific location on the chromosomes of the tomato genome. Cf-4 and Cf-9 are allelic and are located on the short arm of chromosome 1, within a cluster of *HOMOLOGUES OF CLADOSPORIUM RESISTANCE GENE CF-9* (*Hcr9s*) (Jones et al., 1993). Cf-4 is an LRR-RLP containing 25 extracellular LRRs and recognizes Avr4, which is a chitin-binding protein that protects against secreted plant chitinases (Joosten et al., 1994; Thomas et al., 1997; Westerink et al., 2002). Cf-9 contains 27 LRRs in its ectodomain and is only divergent from Cf-4 at its N-terminal part. Cf-9 recognizes Avr9, which is a small cysteine-knotted peptide of unknown function (van den Ackerveken et al., 1992; Jones et al., 1994). Cf-2 and Cf-5 are located on the short arm of chromosome 6, within the *Hcr2* cluster. Cf-2 encodes an LRR-RLP with 38 extracellular LRRs, and recognizes the REQUIRED FOR CLADOSPORIUM RESISTANCE 3 (Rcr3)-Avr2 complex, in which Rcr3 is an apoplastic protease of tomato that is targeted by the secreted protease inhibitor Avr2 of *F. fulva* (Dixon et al., 1996; Dixon et al., 2000; Rooney et al., 2005). Cf-5 is allelic to Cf-2, and is structurally very similar to Cf-2, with 32 extracellular LRRs. Cf-5 matches Avr5, whose function remains unknown (Mesarich et al., 2014; Thomas et al., 1998). Notably, Cf-4 and Cf-9 trigger a much stronger and faster cell death when compared to Cf-2 and Cf-5, but all these Cf proteins confer resistance to *F. fulva*.

Cf-4 is one of the most studied Cf resistance proteins, and several components that are required for Cf-4-mediated immune responses have been identified. SOBIR1 is essential for initiating Cf-4-mediated immune responses such as the HR-related cell death, the apoplastic ROS burst, and MAPK activation (Huang et al., 2021; Liebrand et al., 2013). Moreover, BAK1 is an essential co-receptor required for the Cf-4-triggered apoplastic ROS burst and HR-related cell death (van der Burgh et al., 2019; Postma et al., 2016). Downstream of receptor activation, the helper NB-LRR REQUIRED FOR HR-ASSOCIATED CELL DEATH 3 (NRC3) is required for Cf-4-mediated HR-related cell death (Kourelis et al., 2022), but its contribution to the Cf-mediated apoplastic ROS burst remains unclear. Furthermore, several receptor-like cytoplasmic kinases (RLCKs) have been found to play a role in immune signaling downstream of Cf-4 (Huang et al., 2024). For the functionality of RLP23, which is an *Arabidopsis thaliana* LRR-RLP that recognizes necrosis- and ethylene-inducing-like proteins (NLPs) from bacteria, fungi, and oomycetes, the ENHANCED DISEASE SUSCEPTIBLITY 1 (EDS1)-PHYTOALEXIN DEFICIENT 4 (PAD4) signaling module is essential (Pruitt et al., 2021). However, Cf-4-mediated immunity does not require this EDS1-PAD4 module (Zönnchen et al., 2022). Despite these detailed studies on the molecular background of Cf-4-mediated immunity, the molecular details of Cf-5-mediated immune signaling remain largely unknown. Consequently, the requirements, similarities, and possible differences between Cf-4- and Cf-5-triggered immune responses, except for their differences in intensity of HR-related cell death and speed, are still unknown.

In this study, we aimed to understand whether differences in the intensity of HR-related cell death between Cf-4 and Cf-5 reflect some kind of divergence in their underlying immune signaling mechanisms and the actual level of resistance these Cf proteins provide. Using purified Avr4 and Avr5 proteins in tomato, we confirmed that the Cf-4/Avr4 combination triggers a stronger cell death than the Cf-5/Avr5 combination. Both receptor–effector pairs induced a rapid MAPK activation, whereas the Cf-5/Avr5 combination elicited a more sustained MAPK signaling at later time points. In addition, Avr4 and Avr5 also triggered an apoplastic ROS burst independently of their matching receptors, while remaining dependent on the NADPH oxidase RESPIRATORY BURST OXIDASE HOMOLOG B (RBOHB) for this. Transcriptome analysis revealed that Cf-4 and Cf-5 share a large core set of differentially expressed genes upon the initiation of immune signaling after perception of their matching ligand, but also display qualitative and quantitative differences, with the Cf-5/Avr5 combination inducing a broader and more sustained transcriptional reprogramming. Despite these differences, both Cf-4 and Cf-5 conferred comparable levels of resistance to *F. fulva*. Together, these findings indicate that the intensity of the HR-related cell death does not correlate with the MAPK activation dynamics, apoplastic ROS production, transcriptional reprogramming, or disease resistance.

## Results

### The Cf-5/Avr5-triggered HR-related cell death and apoplastic ROS burst shares the same immune-related signaling pathway with Cf-4/Avr4 in *N. benthamiana*

The Cf-4/Avr4-triggered cell death is a well-studied model when compared to the Cf-5/Avr5-triggered cell death. Concerning Cf-4/Avr4-triggered immunity, it has been shown that SOBIR1, BAK1, and NRC3 are essential for mounting an HR-related cell death (Kourelis et al., 2022; Liebrand et al., 2013; Postma et al., 2016). To determine whether these important immune-related proteins are also required for the Cf-5/Avr5-triggered cell death, cell death assays were performed on mutant lines of *Nicotiana benthamiana* lacking the genes encoding these important immunity-related proteins. For this, the Cf-5/Avr5 combination was transiently expressed in leaves of *N. benthamiana* wild-type (WT)*, sobir1/sobir1-like* (Huang et al., 2021), *bak1* (Zönnchen et al., 2022) and *nrc2/3_3.3.1* (Kourelis et al., 2022) knock-out plants. Cf-4/Avr4 was included as a control in this essay. At 7 dpi, the infiltrated leaves were imaged using red-light imaging, and the intensity of the HR-related cell death was quantified. The results indicate that the Cf-5/Avr5-triggered cell death also requires SOBIR1, BAK1, and NRC2/3, similar to the Cf-4/Avr4-triggered cell death (Figure 1a, b). Moreover, it has been shown that EDS1 and PAD4 are required for RLP23 to trigger an immune response (Pruitt et al., 2021), whereas for Cf-4 these proteins are not required for triggering cell death (Zönnchen et al., 2022). Similar to the Cf-4/Avr4-triggered cell death, EDS1 and PAD4 were also found not to be required for the Cf-5/Avr5-triggered cell death (Figure S1).

**Figure 1:**
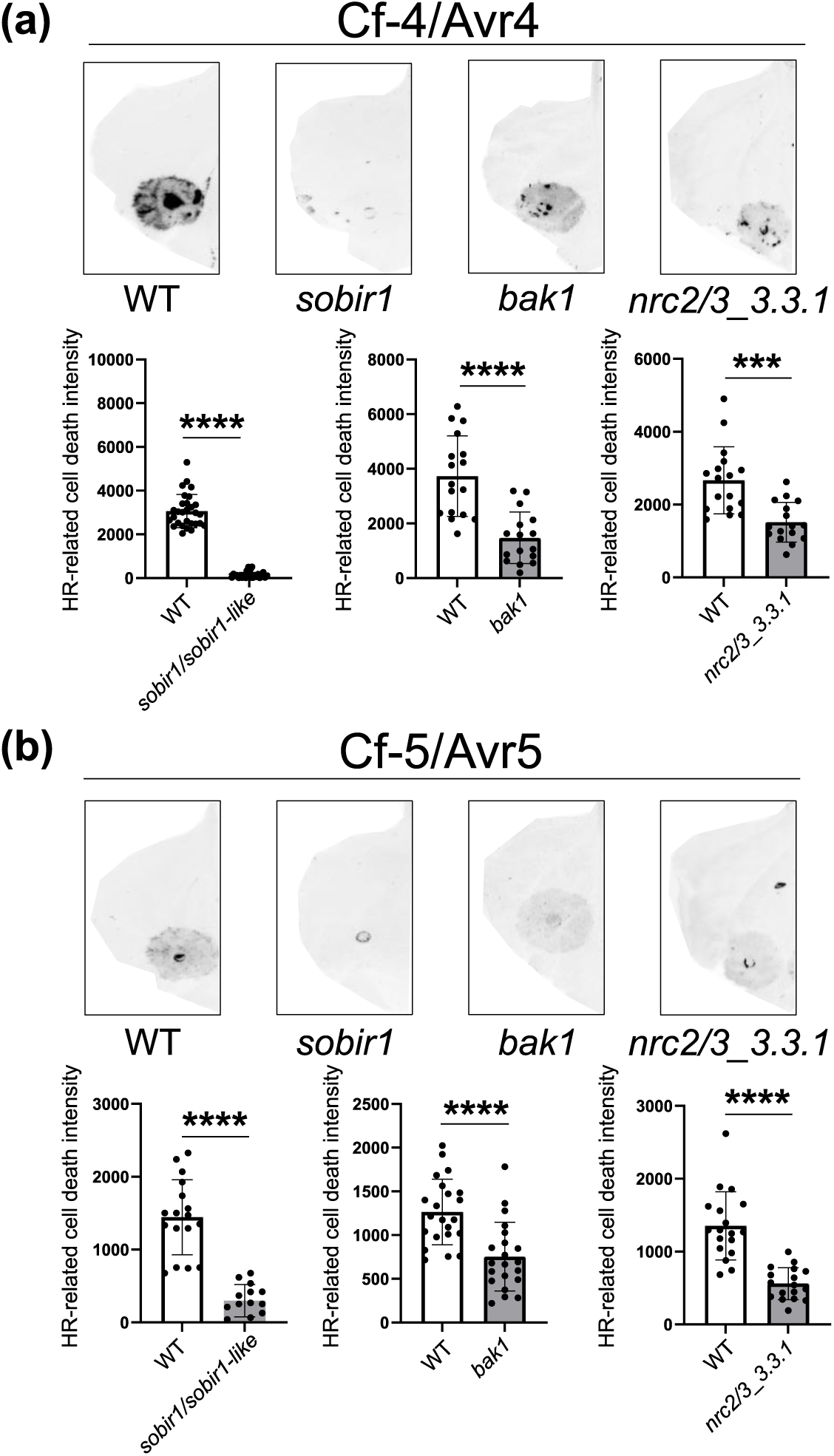
SOBIR1, BAK1 and NRC2/3 are required for Cf-4/Avr4- and Cf-5/Avr5-triggered HR-related cell death. **(a)** Cf-4/Avr4 and **(b)** Cf-5/Avr5 were agroinfiltrated in leaves of *N. benthamiana* wild-type (WT), and the *sobir1/sobir1-like*, *bak1*, and *nrc2/3_3.3.1* knock-out *N. benthamiana* lines (OD_600_ = 0.8). **(a-b)** At 7 dpi, the infiltrated leaves were imaged using red-light imaging, after which the intensity of the HR-related cell death was quantified using ImageLab. The error bars indicate the standard deviation (SD). Statistical significance was determined using a Student *t*-test (unpaired two-tailed *t*-test, p < 0.05). Each mutant was compared to the WT control. The experiments were repeated three times, and representative leaf images are shown. **** p < 0.0001, *** p < 0.001.

The apoplastic ROS burst is also one of the earliest immune responses that is triggered upon Cf-4 activation by its matching ligand (Huang and Joosten, 2025). To assess whether the Cf-4/Avr4- and Cf-5/Avr5-triggered ROS burst also depends on these immune-related proteins, apoplastic ROS burst assays were also performed in the various mutant lines of *N. benthamiana*. For this Cf-4 and Cf-5 were transiently expressed in leaves of these mutant lines, and at 2 dpi, leaf discs taken from the agroinfiltrated leaf areas were challenged with Avr4 and Avr5 protein, respectively, over a period of 5 hours. In the *sobir1/sobir1-like* knock-out plants, the Cf-4/Avr4- and Cf-5/Avr5-triggered ROS was strongly reduced when compared to the WT plants. However, no clear reduction in the apoplastic ROS burst was observed in the *bak1* and *nrc2/3_3.3.1* knock-out plants (Figure 2a, b). Total photon counts were calculated to quantify the overall intensity of the apoplastic ROS burst and to assess whether there were significant differences between the knock-out mutant and WT plants. Only the *sobir1/sobir1-like* mutant showed a significant difference (Figure S2a, b). Previous studies employing a stable transgenic *N. benthamiana* line expressing *Cf-4* had already revealed that SOBIR1 is essential for the Cf-4/Avr4-triggered ROS (Huang et al., 2021), and the present results show that Cf-5 also needs SOBIR1 for the initiation of downstream immune signaling. In contrast, earlier studies using the *bak1* mutant line showed a slightly reduced Cf-4-triggered ROS (Zönnchen et al., 2022). Notably, the results of the ROS burst assays in the *nrc2/3_3.3.1* mutant plants indicated that the Cf-4- and Cf-5-triggered ROS burst is NRC2- and NRC3-independent. Finally, both Cf-4 and Cf-5 triggered a biphasic ROS burst in *N. benthamiana*, whereas Cf-9 also triggered a biphasic ROS burst upon challenge with Avr9 (Figure S3). Furthermore, the total photon counts displayed no significant differences in the overall ROS intensity when the Cf-4-, Cf-5-, and Cf-9-triggered ROS were compared (Figure S3).

**Figure 2:**
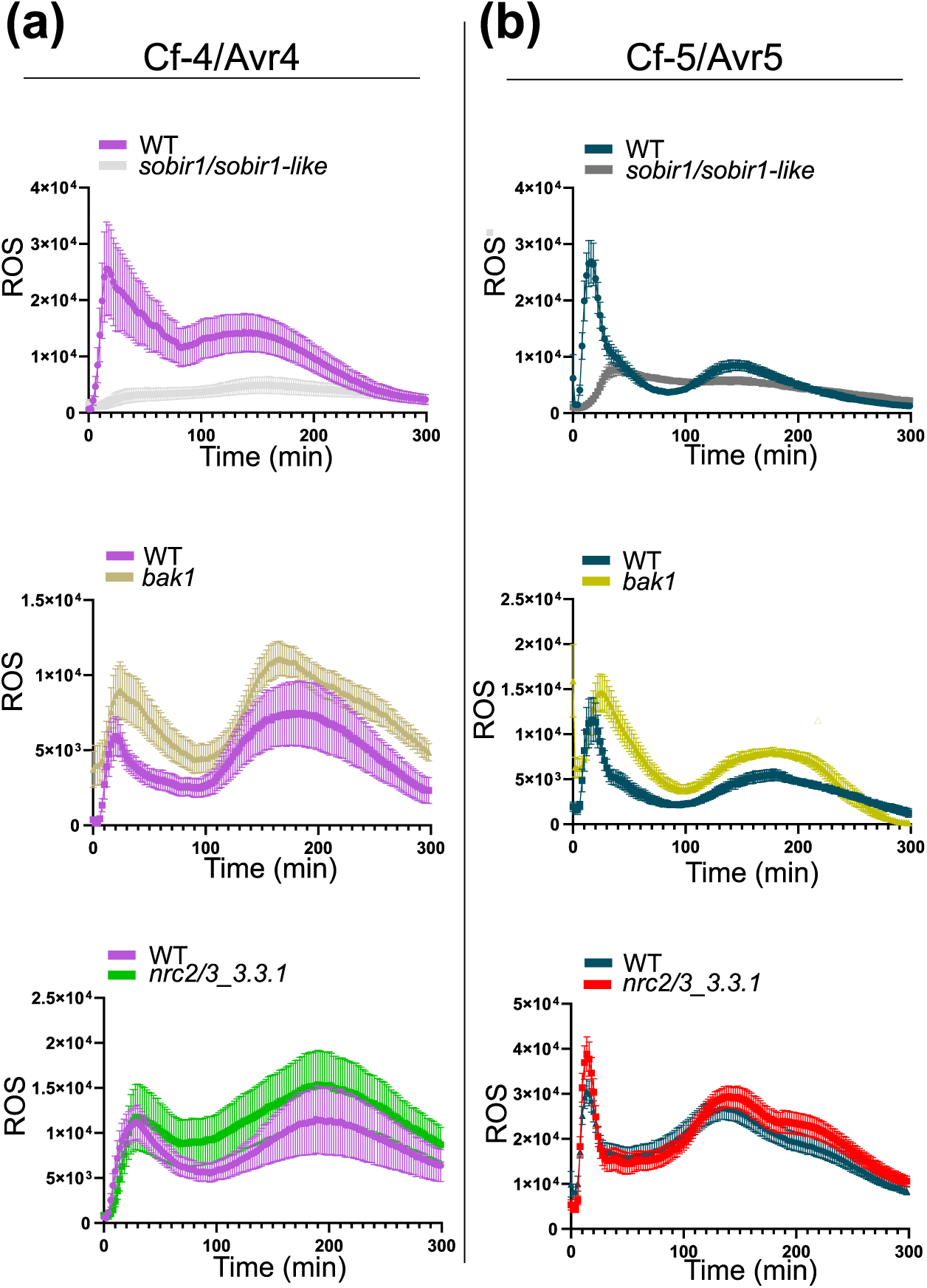
The *sobir1/sobir1-like* mutant plants show a strong reduction in the Cf-4- and Cf-5-triggered ROS burst, whereas the *bak1* and *nrc2/3_3.3.1* mutants do not. Cf-4 **(a)** and Cf-5 **(b)** were transiently expressed in leaves of *N. benthamiana* wild-type (WT), and the *sobir1/sobir1-like*, *bak1*, and *nrc2/3_3.3.1* knock-out *N. benthamiana* lines (OD_600_ = 0.1). At 2 dpi, leaf discs taken from infiltrated leaf areas were challenged with 0.1 µM Avr4 protein **(a)** and 0.1 µM Avr5 protein **(b)** over a period of 5 hours. The apoplastic ROS burst was measured using a luminol-based assay. Error bars indicate the standard error of the mean (SEM). The experiment was repeated at least three times, and representative results are shown.

### Cf-4/Avr4 and Cf-5/Avr5 differ in the intensity of the HR-related cell death that is triggered and in MAPK activation dynamics

Previous work showed that the Cf-4/Avr4 combination triggers a stronger and faster cell death when compared to the Cf-5/Avr5 combination, both in *N. benthamiana* and tomato (Chapter 2). Here, we tested whether this difference is also observed when using pure Avr4 and Avr5 proteins instead of apoplastic fluid containing the Avrs. For the HR-related cell death assay, 5 µM solutions of Avr4 and Avr5, or milli-Q water (MQ), as a control, were infiltrated into leaves of MM:Cf-4, MM:Cf-5 and MM:Cf-0 tomato plants, after which the intensity of HR-related cell death was imaged at 7 dpi by red-light imaging. Again, Avr4 triggered a stronger and faster cell death in leaves of MM:Cf-4 plants than Avr5 does in leaves of MM:Cf-5 tomato (Figure 3a). As expected, no HR-related cell death was observed in non-matching Cf receptor-Avr effector combinations, and neither Avr4 nor Avr5 triggered an HR-related cell death in leaves of MM:Cf-0 plants, which showed responses similar to the MQ control (Figure 3a). The results show that the differences in strength of cell death and timing, using pure Avr4 and Avr5 proteins are consistent with the previously reported study, which assessed the strength of cell death using apoplastic fluid containing Avr4 and Avr5.

**Figure 3:**
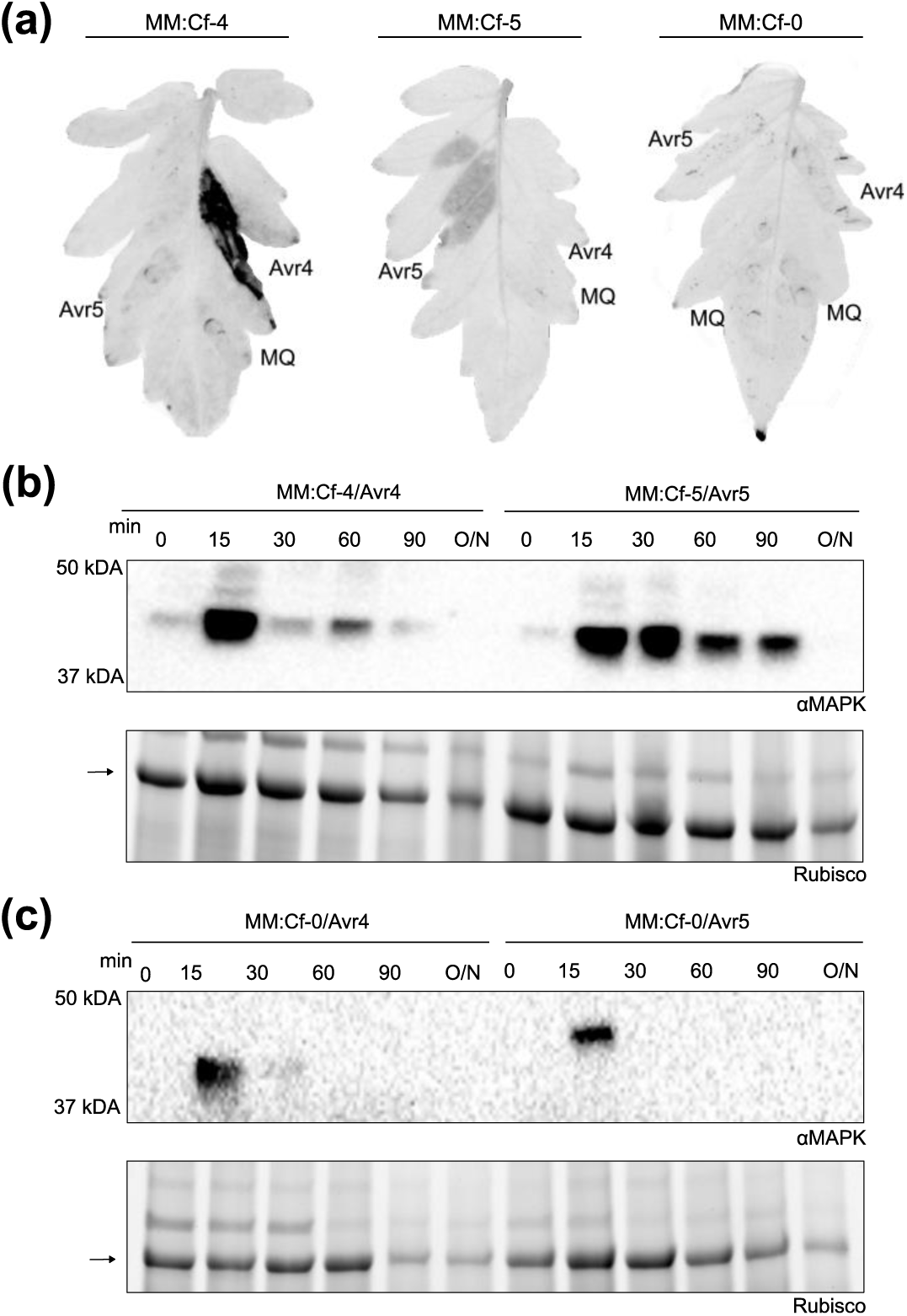
The Cf-4/Avr4 combination triggers a strong HR, whereas Cf-5/Avr5 triggers a stronger, more sustained MAPK activation in tomato, when compared to Cf-4/Avr4. **(a)** Solutions of 5 µM of Avr4 and Avr5, and MQ as a negative control, were infiltrated in leaves of MM:Cf-4, MM:Cf-5, and MM:Cf-0 tomato plants. At 7 dpi, the leaves were imaged using red-light imaging. The experiment was repeated three times and representative images are shown. **(b)** Solutions of 5 µM Avr4 and Avr5 were infiltrated in leaves of MM:Cf-4 and MM:Cf-5 plants, respectively. **(c)** Solutions of 5 µM Avr4 and Avr5 were infiltrated in leaves of MM:Cf-0 plants. **(b-c)** Infiltrated leaves were harvested at 0, 15, 30, 60, 90 minutes post infiltration and after incubation overnight (O/N), after which total proteins were extracted and subjected to immunoblotting using phosphorylated 44/42 MAPK antibody to detect the activated MAPKs. The amount of total protein that was loaded was determined by using the intensity of the Rubisco band (see black arrow on the left) as a reference. Note that the exposure time in C (upper panel, 70 sec) is much longer than the exposure time in B (upper panel, 13 sec). The experiment was repeated three times for **(b)**, and two times for **(c)**, and representative results are shown.

Swift MAPK activation is one of the robust immune responses triggered by a matching Cf/Avr combination (Huang, 2022; Huang et al., 2024; Huang and Joosten, 2025). To assess whether the MAPK activation dynamics differ between the Cf-4/Avr4- and Cf-5/Avr5-triggered immune responses, we performed MAPK activation assays at different time points after challenge with the matching Avr proteins and compared the activation levels of the MAPKs after 15 minutes, which was earlier found to be the time point of maximum MAPK activation for the Cf-4/Avr4 combination (Huang, 2022). This approach also allowed us to evaluate whether differences in the intensity of MAPK activation or dynamics might correlate with differences in the strength of cell death between Cf-4/Avr4 and Cf-5/Avr5. For this, 5 µM solutions of Avr4 and Avr5 were again infiltrated in leaves of MM:Cf-4 and MM:Cf-5 tomato plants, and as a negative control, they were also infiltrated in leaves of MM:Cf-0 plants. Furthermore, a 5 µM solution of the flg22 peptide, matching endogenous FLS2, was also included. The infiltrated leaves were harvested at 0, 15, 30, 60, 90 mins, and after overnight (O/N) incubation. At 15 minutes, both the Cf-4/Avr4 and Cf-5/Avr5 treatments resulted in a strong MAPK activation (Figure 3b). After normalization to the intensity of the corresponding Rubisco band that was used as a total protein loading control, no difference in the strength of MAPK activation was detected between Cf-4/Avr4 and Cf-5/Avr5 after a 15-minute challenge (Figure S4a). At later time points (30, 60, and 90 minutes) the Cf-5/Avr5 combination had triggered a stronger and more sustained MAPK activation as compared to Cf-4/Avr4 (Figure 3b). In leaves of MM:Cf-0 plants, only weak activation of MAPKs was detected at 15 minutes (Figure 3c). Moreover, the infiltration of flg22 induced strong MAPK activation at 15 minutes, which rapidly decreased at later time points (Figure S4b).

These results indicate that both Cf-4/Avr4 and Cf-5/Avr5 induced MAPK activation with a similar intensity at 15 minutes after challenge. However, the Cf-5/Avr5 combination resulted in a stronger and more sustained MAPK activation at later time points. Moreover, the Cf/Avr-triggered MAPK activation was more sustained than the flg22-triggered MAPK activation.

### In leaves of tomato, Avr4 and Avr5 trigger an apoplastic ROS burst also in the absence of their matching receptor

To assess whether the intensity and profile of the apoplastic ROS burst differ between the Cf-4/Avr4- and Cf-5/Avr5-triggered immune responses in tomato, we conducted apoplastic ROS burst assays employing the protocol previously used in *N. benthamiana.* First, we tested the method by challenging MM:Cf-0, tomato cultivar Rio Grande, and a Rio Grande *rbohb* mutant that were obtained from the Biotechnology Center at the Boyce Thompson Institute (Leuschen-Kohl et al., 2025), with flg22. Leaf discs were challenged with a 0.1 µM solution of flg22 over a period of 5 hours, which for MM:Cf-0 resulted in a swift monophasic ROS burst, whereas a MQ control did not trigger any response (Figure S5a). Similarly, the flg22-triggered ROS burst was monophasic and fast in Rio Grande, whereas no response was detected in the Rio Grande *rbohb* mutant plants (Figure S5b). These results indicate that the assay conditions reliably detect the flg22-triggered ROS burst in tomato, and are suitable for comparing the Cf-4/Avr4- and Cf-5/Avr5-triggered a ROS burst. Furthermore, this observation proves that RBOHB from tomato is solely responsible for the apoplastic ROS burst upon challenge with an ExIP.

For the Cf-4/Avr4- and Cf-5/Avr5-triggered ROS burst assays, leaf discs were taken from MM:Cf-4 and MM:Cf-5 tomato plants, and challenged with a 1µM solution of Avr4 and Avr5, respectively. Again, MQ was used as a negative control for both genotypes. Both the Cf-4/Avr4 and Cf-5/Avr5 combination resulted in a ROS burst profile that was clearly distinct from the MQ control. In leaf discs taken from MM:Cf-4 plants, challenge with Avr4 induced a biphasic ROS burst. However, a late increment in the ROS profile was also observed upon treatment with MQ (Figure 4a). Notably, the biphasic ROS profile of the MM:Cf-4/Avr4 response was not fully consistent between experiments, and in some assays, the Cf-4/Avr4 combination triggered a broader, monophasic ROS burst. In leaf discs taken from MM:Cf-5 plants, Avr5 also triggered a broad, monophasic ROS burst, although a late increase in ROS was also detected in the MQ control (Figure 4a). Total photon counts were calculated to assess the intensity of the ROS burst, now also including the Cf-9/Avr9 combination. No differences in total photon counts were detected between Cf-4/Avr4 and Cf-5/Avr5. In contrast, challenge of leaf discs of Cf-9 tomato plants with a 1 µM solution of Avr9 resulted in a less intense ROS burst than the Cf-4/Avr4 and Cf-5/Avr5 combinations (Figure S6). Because the Avr9-triggered ROS burst was not clearly detectable, a challenge with Avr9 was also performed with the *Hcr9-9D* tomato line, which expresses only the *Cf- 9* gene. However, it still remained difficult to observe a clear Avr9-induced ROS burst compared to MQ control, both in leaf discs from *Hcr9-9D* and MM:Cf-9 plants (Figure S7a, b). Moreover, leaf discs of MM:Cf-4, MM:Cf-5 and MM:Cf-9 plants were also challenged with Avr4 and Avr5 to assess responses of non-matching receptor-effector combinations. MM:Cf-0 was taken along as a negative control. Unexpectedly, both Avr4 and Avr5 triggered a ROS burst in all lines (Figure 4b), with no obvious differences in intensity among the different lines (Figure S8a, b). Lastly, Rio Grande and the Rio Grande *rbohb* knock-out line were included to test whether Avr4 and Avr5 trigger a ROS burst that is RBOHB-dependent. Both Avr4 and Avr5 triggered a ROS burst in Rio Grande, whereas no ROS burst was detected in the *rbohb* mutant lines (Figure S9).

**Figure 4:**
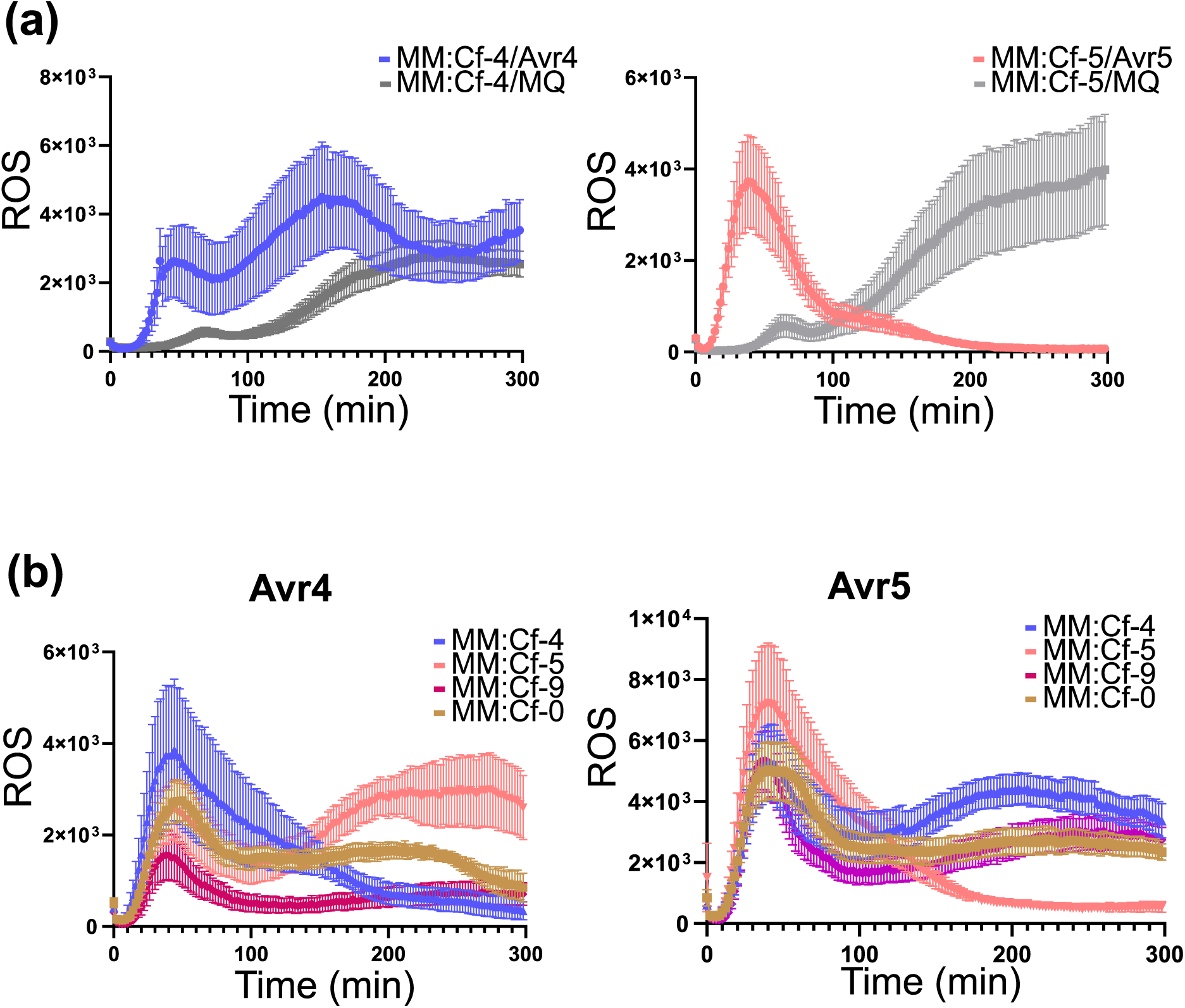
Avr4 and Avr5 both induce a ROS burst in leaves of MM:Cf-4, MM:Cf-5, MM:Cf-9 and MM:Cf-0 tomato. **(a)** Leaf discs from MM:Cf-4 and MM:Cf-5 were challenged with a 1 µM solution of Avr4 and Avr5, respectively, over a period of 5 hours. MQ served as a negative control. **(b)** Leaf discs of leaves of MM:Cf-4, MM:Cf-5, MM:Cf-9, and MM:Cf-0 plants were challenged with a 1 µM solution of Avr4 and Avr5. **(a-b)** The apoplastic ROS burst was measured using a luminol-based assay. Error bars show the SEM.

Overall, these results indicate that Avr4 and Avr5 both trigger a ROS burst of similar intensity in leaf discs of tomato plants expressing their matching receptor. Interestingly, both Avr4 and Avr5 also trigger a ROS burst in non-matching receptor/effector combinations, and also in tomato MM-Cf0 and Rio Grande. Furthermore, the Avr4- and Avr5-triggered ROS burst was found to be fully dependent on RBOHB. Although Avr9 triggers a ROS burst in *N. benthamiana* transiently expressing *Cf-9* (Figure S3), we did not detect a clear Avr9-induced ROS burst in MM-Cf-9 plants and the additional tomato lines that were tested.

### The matching Cf-4/Avr4 and Cf-5/Avr5 combinations trigger quantitative and qualitative differences in transcriptional reprogramming in tomato

To assess differences and similarities between Cf-4/Avr4- and Cf-5/Avr5-triggered immunity, we conducted RNA-seq experiments at 3h and 7h after infiltration of leaflets of MM-Cf-4 and MM-Cf-5 tomato plants with a solution of 0,5 µM of Avr4 and Avr5 protein, respectively. The overlap of the differentially expressed genes (DEGs), identified relative to their respective 0h time point, between the various treatments and time points was visualized using hierarchical clustering heatmap and Venn diagrams. At 3h after infiltration, the Cf-4/Avr4 combination (A) resulted in 8,706 DEGs, whereas the Cf-5/Avr5 combination (B) resulted in a higher number of DEGs, being 11,123, with 7,463 DEGs being shared between the two treatments (Figure 5). At 7h after infiltration, the total number of DEGs decreased for both treatments, as challenge of Cf-4 with Avr4 (C) resulted in 5,141 DEGs, whereas challenge of Cf-5 with Avr5 (D) resulted in 7,576 DEGs, of which 3,522 DEGs were shared (Figure 5). Comparison of all four treatments revealed a large set of shared DEGs (2553), indicating that both immune pathways share a core transcriptional response (Figure 5). However, each treatment also showed a distinct set of DEGs. Notably, the Cf-5/Avr5 combination consistently resulted in a higher number of DEGs at both time points when compared to Cf-4/Avr4, suggesting that the Cf-5/Avr5-triggered immune signaling pathway overall induces a broader transcriptional reprogramming than immune signaling triggered by Cf-4 upon perception of Avr4 (Figure 5).

**Figure 5:**
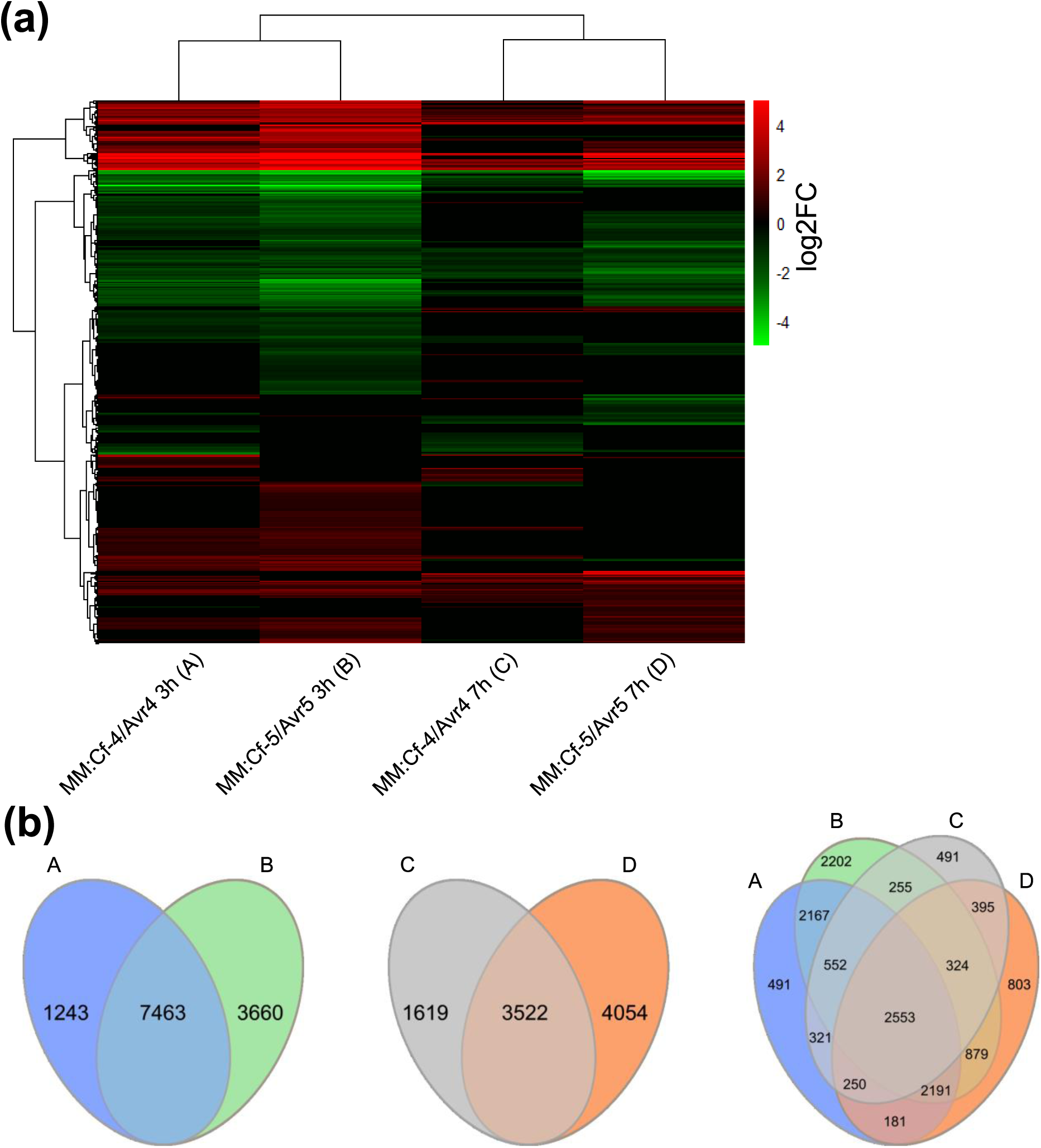
Comparison of the overall transcriptional regulation between the Cf-4/Avr4 and Cf-5/Avr5 combinations, at 3h and 7h after challenge. **(a)** Hierarchical clustering heatmap of differentially expressed genes (DEGs) across the indicated treatments. Gene expression values are shown as log2 fold change (log2FC), with red indicating upregulation, green indicating downregulation, and black indicating no significant change in gene transcription. **(b)** Overlap and differences in DEGs at 3h and at 7h, between Cf-4/Avr4 and Cf-5/Avr5, are visualized using Venn diagrams. A, B, C and D refer to the treatments specified in **(a)**.

Phytohormones are key components of plant defense (Denancé et al., 2013). Therefore, we assessed whether phytohormone-related genes were differentially transcriptionally regulated between the various treatments (Figure 6a). For salicylic acid (SA)-related gene regulation, the transcriptional regulation of the genes encoding NONEXPRESSOR OF PR GENES1 (NPR1) and basic leucine zipper (bZIP) TGA element-binding transcription factors (TFs) was studied. Among the TGA family of TFs, *Solyc12g056860* and *Solyc10g080770* were both moderately upregulated at the 3h time point in both treatments, with a stronger induction in the Cf-5/Avr5 combination. At 7h, the expression of *Solyc12g056860* was sustained only in Cf-4/Avr4, while *Solyc10g080770* remained upregulated only in Cf-5/Avr5. *Solyc11g064950*, *Solyc04g011670*, and *Solyc10g080780*, which all also encode TGA-TFs, were only significantly differentially regulated in Cf-5/Avr5 at 3h, with *Solyc11g064950* being upregulated while *Solyc04g011670* and *Solyc10g080780* were downregulated. In contrast, the genes encoding the Solyc05g009660 and Solyc06g074320 TGA-TFs were downregulated across all conditions and time points. Solyc10g080770, which encodes NPR1, the master regulator of SA-related immune responses, was upregulated at 3h in both treatments and remained differentially expressed in Cf-5/Avr5 at 7h. Similarly, two genes encoding NPR1-INTERACTING (NIM1-INTERACTING) proteins, *Solyc02g069310* and *Solyc07g044980*, were upregulated at 3h in both immune pathways, but at 7h their differential expression was maintained only in Cf-5/Avr5 (Figure 6a).

**Figure 6:**
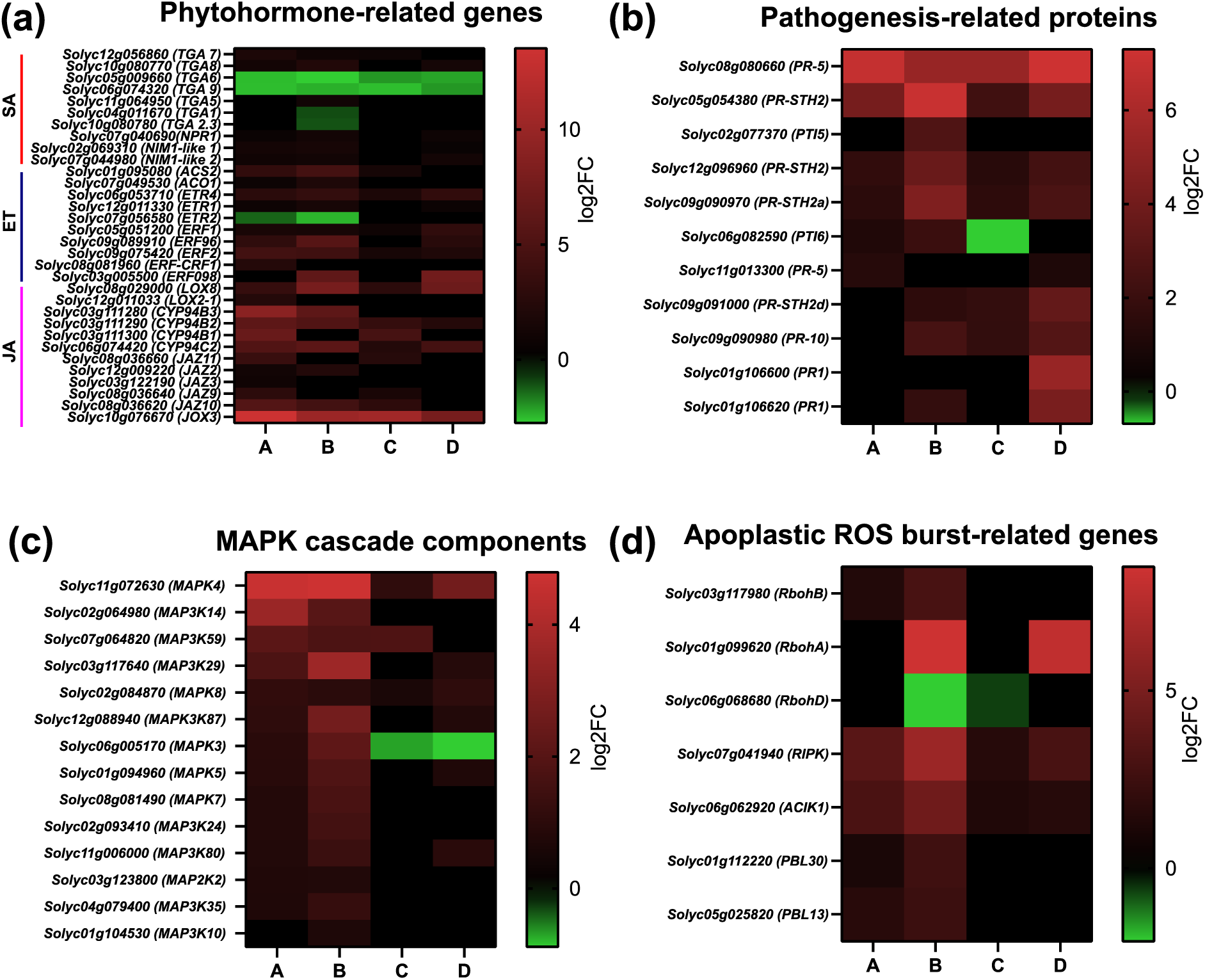
Shared and distinct transcriptional regulation of immune-related genes triggered by Cf-4/Avr4 and Cf-5/Avr5 in tomato. Heatmaps show the expression levels of selected genes involved in **(a)** phytohormone signaling, **(b)** encoding pathogenesis-related proteins and **(c)** MAPK cascade components, and **(d)** genes involved in the apoplastic ROS burst, for the Cf-4/Avr4 and Cf-5/Avr5 combination. A, Cf-4/Avr4 at 3h; B, Cf-5/Avr5 at 3h; C, Cf-4/Avr4 at 7h; D, Cf-5/Avr5 at 7h.

Concerning ethylene (ET)-related signaling, *Solyc01g095080*, which encodes 1-AMINOCYCLOPROPANE-1-CARBOXYLIC ACID (ACC) SYNTHASE 2 (ACS2), which is the key enzyme for ethylene biosynthesis, was upregulated at 3h in both immune pathways, with consistently higher expression in Cf-5/Avr5 than in Cf-4/Avr4. Similarly, ACS2 was also upregulated upon *F. fulva* race 5 inoculation of resistant MM-Cf-4 tomato, as well as in dying seedlings expressing both Cf-4 and Avr4 when transferred to 20°C, thereby inducing the defense response (Etalo et al., 2013). At 7h, *ACS2* showed lower expression in Cf-4/Avr4 (log2FC 1,4) compared Cf-4/Avr4 at 3h), and the gene was not differentially regulated anymore at 7h in the Cf-5/Avr5 combination. *Solyc07g049530*, encoding 1- AMINOCYCLOPROPANE-1-CARBOXYLATE OXIDASE 1 (ACO1), which is involved in the ethylene production, showed modest upregulation at 3h, but was not differentially expressed at 7h in either combination. In ethylene perception, Solyc06g053710 (ETHYLENE RECEPTOR 4 (ETR4)) and Solyc12g011330 (ETR1) play a role, and the encoding genes were upregulated at 3h in both combinations. Notably, ETR4 remained strongly upregulated at 7h, particularly in Cf-5/Avr5, whereas the upregulation of ETR1 expression was transient. In contrast, *Solyc07g056580* (*ETR2*) was downregulated at 3h. The gene encoding Solyc05g051200 (ETHYLENE-RESPONSIVE TRANSCRIPTION FACTOR 1

(ERF1)) was induced in Cf-4/Avr4 at both time points and showed strong upregulation in the Cf-5/Avr5 combination at 7h (Figure 5b). ERF-encoding genes associated with MAPK signaling and plant-pathogen interaction pathways in the KEGG analysis were selected to be included in the heatmap (Figure 6a). Genes encoding Solyc09g089910 (ERF096) and Solyc09g075420 (ERF2), were strongly induced at 3h for both combinations, with *ERF096* showing particularly high expression levels in Avr5-challenged Cf-5 tomato. *Solyc03g005500* (*ERF098*) was specifically strongly upregulated in Cf-5/Avr5 at both time-points, whereas *Solyc08g081960* (encoding the ERF referred to as CYTOKININ RESPONSE FACTOR 11 (CRF11)) was significantly upregulated only in Cf-4/Avr4 at 3h (Figure 6a).

Concerning jasmonic acid (JA)-related signaling, *Solyc08g029000*, encoding LIPOXYGENASE 8 (LOX8), which is involved in JA synthesis, was up-regulated in both treatments at 3h and 7h, but showed higher and more sustained expression in Cf-5/Avr5. In contrast, *Solyc12g011033* (*LOX2-1*) was specifically upregulated in Cf-4/Avr4 at 3h. Among the CYTOCHROME P450 (CYP) subfamily of the CYP94s, which are involved in JA degradation, *Solyc03g111290* (*CYP94B2*) and *Solyc06g074420* (*CYP94C2*) were strongly upregulated in both treatments at 3h and 7h. *Solyc03g111280* (*CYP94B3*) was also upregulated in both treatments, but only at 3h, with stronger upregulation in Cf-4/Avr4. *Solyc03g111300* (*CYP94B1*) was exclusively upregulated in Cf-4/Avr4 at both time points. The gene encoding Solyc10g011650 (JASMONATE RESPONSE 1 (JAR1)-like) was upregulated only in Cf-4/Avr4 at 3h. Solyc10g076670 (JASMONATE-INDUCED OXYGENASE 3 (JOX3)), which degrades an excess of JA and serves as a marker for elevated JA levels (Caarls et al., 2017), was extremely highly upregulated in both combinations at both time points. Genes encoding several negative regulators of JA signaling, including *Solyc08g036620* (*JASMONATE ZIM-DOMAIN PROTEIN 10* (*JAZ10*)) were induced at 3h in both treatments, and remained significantly upregulated at 7h in the Cf-4/Avr4 combination. *Solyc08g036660* (*JAZ11*) and *Solyc08g036640* (*JAZ9*) were significantly upregulated only in Cf-4/Avr4 at 3h and 7h (Figure 6a).

Pathogenesis-related proteins (PRs) are key proteins of plant immunity, functioning either as direct antimicrobials or as regulators of defense signaling (van Loon et al., 2006; Sels et al., 2008). *Solyc08g08066*, the gene encoding PR-5, which is a thaumatin-like protein, showed strong upregulation in both treatments, both at 3h and 7h. PR-5 has been shown to enhance resistance against pathogens, specifically by stimulation of β-1,3-glucanase activity (Li et al., 2023). *Solyc05g054380* (encoding SALT TOLERANCE HOMOLOG 2 (PR-STH2)), was strongly upregulated in Cf-5/Avr5. Furthermore, the genes encoding Solyc09g091000 (PR-STH2d) andSolyc09g090970 (PR-STH2a), which are interactors of *Sl*WRKY30 (Dang et al., 2023), and Solyc02g065470, encoding a PR-1 like protein, were upregulated at both time points for the Cf-4/Avr4 and Cf-5/Avr5 combination. In contrast, some PR genes showed time-and Cf/Avr-specific regulation. For example, *Solyc09g091000* (*PR-STH2d*) and *Solyc09g090980* (*PR-10*) were upregulated in both combinations at 7h, but at 3h, these genes were only significantly upregulated in the Cf-5/Avr5 combination. Solyc06g082590 (PTO-INTERACTOR 6 (PTI6)) and Solyc02g077370 (PTI5) are transcription factors known to activate defense-related gene expression (Gu et al., 2002). *PTI6* showed early induction in both treatments at 3h, but was eventually downregulated in Cf-4/Avr4 and was non-responsive in Cf-5/Avr5 at 7h. *PTI5* was specifically upregulated in the case of the Cf-5/Avr5 combination at 3h. *Solyc01g106620* (*PR1*) was transcriptionally induced at 7h in both pathways, with much stronger expression in Cf-5/Avr5, while *Solyc01g106600* (encoding PR1) was induced only at 7h (Figure 6b).

Mitogen-activated kinases (MAPKs) are also regulators of plant immunity, defense, and phytohormone production. MAPKs have, for example, been shown to play an important role in the resistance response of tomato to *F. fulva* (Stulemeijer et al., 2007). We have assessed the transcriptional regulation of the various *MAPKs* and their upstream *MAPKKs* and *MAPKKKs*. The gene encoding Solyc11g072630 (MAPK4), which has been reported to be induced during ethylene-related processes (Li et al., 2017; Zhang et al., 2024), exhibited the highest upregulation among all MAPK-encoding genes at 3h in both combinations, with stronger induction for Cf-5/Avr5. Although *MAPK4* expression levels decreased at 7h, they remained upregulated in both treatments. *Solyc02g064980*, encoding MAPK3K14, which is involved in the response to salt stress (Meng et al., 2020), showed induction at 3h, with higher expression levels for Cf-4/Avr4. The genes encoding the MAPKKKs, MAPKKs and MAPKs, Solyc03g117640 (MAP3K29), Solyc12g088940 (MAP3K87), Solyc06g005170 (MAPK3), Solyc01g094960 (MAPK5), Solyc08g081490 (MPK7), Solyc02g093410 (MAP3K24), Solyc11g006000 (MAP3K80), Solyc03g123800 (MAPK2K2), and Solyc04g079400 (MAP3K35), were all upregulated at 3h in both combinations, with generally stronger induction in the Cf-5/Avr5 combination, indicating enhanced early MAPK-associated transcriptional activation in this interaction. Notably, *MAPK3* was downregulated at 7h in both treatments, suggesting a negative feedback regulation during later stages of the immune response. At 7h, a subset of MAPK-encoding genes displayed Cf/Avr-specific regulation. *MAP3K87*, *MAPK5*, *MAP3K80*, and *Solyc01g104530* (*MAPK3K10*) remained upregulated exclusively in the Cf-5/Avr5 combination, indicating prolonged MAPK-associated signaling in this Cf/Avr combination (Figure 6c).

Next, we evaluated the transcriptional behavior of apoplastic ROS burst-related genes across the various samples. *Solyc03g117980*, encoding RBOHB, which has been functionally characterized concerning its requirement for the flg22-triggered apoplastic ROS burst in tomato (Li et al., 2015), was upregulated at 3h in both combinations, but showed no regulation at 7h. Interestingly, *Solyc01g099620* (*RBOHA*), which has been found to be upregulated during biotic stress responses (Zhou et al., 2018), was specifically and strongly upregulated only during the Cf-5/Avr5-triggered response at 3h and 7h, while remaining transcriptionally unchanged in the Cf-4/Avr4 samples. We showed that Avr4 and Avr5 trigger an apoplastic ROS burst in leaves of MM:Cf-0 tomato plants, and the control infiltration of Avr4 and Avr5 in leaves of MM:Cf-0 resulted in an upregulation of *RBOHB*, but not *RBOHA*. The results indicate that the transcriptional activation of *RBOHA* is specific for Cf-5/Avr5-triggered immunity (Figure 6d). In *N. benthamiana*, receptor-like cytoplasmic kinase (RLCK) subfamilies VII-6, -7, and -8 have been shown to be required for a full Cf-4/Avr4-triggered apoplastic ROS burst (Huang et al., 2024). In tomato, Solyc07g041940 (RPM1-INDUCED PROTEIN KINASE (RIPK)), which is an RLCK VII-6 member, has been shown to be required for the chitin-triggered ROS burst (Wang et al., 2022), supporting a conserved role for this RLCK VII family in apoplastic ROS signaling. Consistent with these findings and *RBOHB* transcriptional regulation, *RIPK* was strongly upregulated at 3h, with even higher upregulation in Cf-5/Avr5 compared to Cf-4/Avr4, and reduced expression at 7h for both combinations. The genes encoding Solyc06g062920 (AVR9/CF-9 INDUCED KINASE (ACIK1); an RLCK VII-6 member), Solyc01g112220 (AVRPPHB-SUSCEPTIBLE 1 (PBS1)-LIKE 30 (PBL30); an RLCK VII-7 member), and Solyc05g025820 (PBL13/RIPK-like; an RLCK VII-6 member), all showed strong upregulation at 3h. Among these, only *ACIK1* was still expressed at 7h, whereas the other *RLCKs* were not detectably transcriptionally induced at this later time point (Figure 6d).

Moreover, we also focused on the transcriptional regulation of WRKY transcription factors, which are named after their “WRKY” DNA-binding motif and play key roles in transcriptional reprogramming during plant defense (Rushton et al., 2010; Wani et al., 2021). In the Cf-4/Avr4-triggered immune response, 33 *WRKY* genes were differentially regulated at 3h, while 21 *WRKY* genes were differentially regulated at 7h. In contrast, the Cf-5/Avr5 combination triggered an even stronger *WRKY* transcriptional response, with 43 differentially expressed *WRKY* genes at 3h and 29 *WRKY* genes at 7h, although a large proportion of the transcriptionally regulated WRKY transcription factors were overlapping between the two combinations (Figure S10a). *Solyc04g051540* (*WRKY13*) and *Solyc07g051840* (*WRKY17*) were differentially expressed exclusively for the Cf-4/Avr4 combination at both time points. In contrast, *Solyc05g007110* (*WRKY76*) and Solyc05g050340 (*WRKY58*) were specifically regulated only for Cf-5/Avr5 at both 3h and 7h. Furthermore, several *WRKYs* were uniquely differentially expressed in the Cf-5/Avr5 combination at 3h, including *Solyc06g048870* (*WRKY19*), *Solyc03g007380* (*WRKY52*), *Solyc06g068460* (*WRKY40*), *Solyc04g051690* (*WRKY51*), *Solyc09g015770* (*WRKY81*), *Solyc12g014610* (*WRKY20*), and *Solyc07g005650* (*WRKY32*). At 7h, *Solyc09g066010* (*WRKY24*) was uniquely differentially expressed in the Cf-5/Avr5 combination (Figure S10a).

We further analyzed differentially regulated PCD-related genes, including genes encoding metacaspases (MCs), vacuolar processing enzymes (VPEs), and BCL2-ASSOCIATED X PROTEIN (BAX1) inhibitor proteins (BIs). Among the MCs, genes encoding Solyc01g105320 (MC4), Solyc01g105300 (MC5), and Solyc10g081300 (MC8) showed upregulation at 3h for Cf-4/Avr4, with a higher expression level when compared to Cf-5/Avr5 at the same time point. Several other *MCs*, including *Solyc05g052130* (*MC3*), *Solyc09g098150* (*MC7*), *Solyc03g094160* (*MC2*), and *Solyc01g088710* (*MC1*), displayed modest upregulation at 3h in both combinations, with slightly higher expression levels for Cf-5/Avr5 (Figure S10b). *MC3* remained upregulated at 7h, and *MC5* showed a sustained induction at 7h, specifically in the Cf-5/Avr5 combination. In addition, *MC8* was only upregulated at both 3h and 7h in Cf-4/Avr4, whereas *MC6* was upregulated only at 7h in Cf-5/Avr5. Concerning genes encoding VPEs, *Solyc08g065710* (*VPE11*) was strongly induced at 3h in both combinations, with higher expression in Cf-4/Avr4. At 7h, *VPE11* expression remained elevated only in Cf-5/Avr5, while it was not regulated in Cf-4/Avr4. The other *VPEs* showed no strong induction or were downregulated at 3h. The genes encoding BIs showed overall induction in both combinations, with differences primarily in expression levels and duration. *Solyc07g042510*, encoding a BI 1 family protein, was strongly upregulated at 3h in both the Cf-4/Avr4 and Cf-5/Avr5 combination, and its expression increased at 7 h in Cf-5/Avr5, while decreasing in Cf-4/Avr4. *Solyc08g077980*, encoding another BI protein, showed modest induction at 3h and remained upregulated at 7h, only in Cf-5/Avr5. *Solyc09g065610*, also encoding a BI, was induced at 7h in both combinations, with a higher expression level in Cf-5/Avr5, and showed induction at 3h, exclusively in Cf-5/Avr5 (Figure S10b).

### Both Cf-4 and Cf-5 confer robust resistance to *F. fulva*

To determine whether the distinct immune responses between Cf-4/Avr4 and Cf-5/Avr5 affect the level of actual resistance against *F. fulva*, we performed *F. fulva* race 0 inoculation assays on MM:Cf-4 and MM:Cf-5 tomato plants. Tomato MM:Cf-0 was used as the susceptible control. Race 0 produces all known effectors, including Avr4 and Avr5, and is therefore avirulent on tomato carrying the Cf-4 or Cf-5 gene. Inoculated leaves were harvested at 4, 7, 11, and 15 dpi, followed by genomic DNA extraction to quantify the fungal biomass by qPCR.

MM:Cf-0 showed a significantly higher fungal biomass at 7, 11, and 15 dpi, when compared to MM:Cf-4 and MM:Cf-5, whereas no significant differences were detected among the latter two genotypes. Importantly, when comparing MM:Cf-4 and MM:Cf-5, no significant differences in fungal biomass were observed at any time point. The results indicate that both Cf-4 and Cf-5 provide robust resistance to *F. fulva* race 0 (Figure 7a).

**Figure 7:**
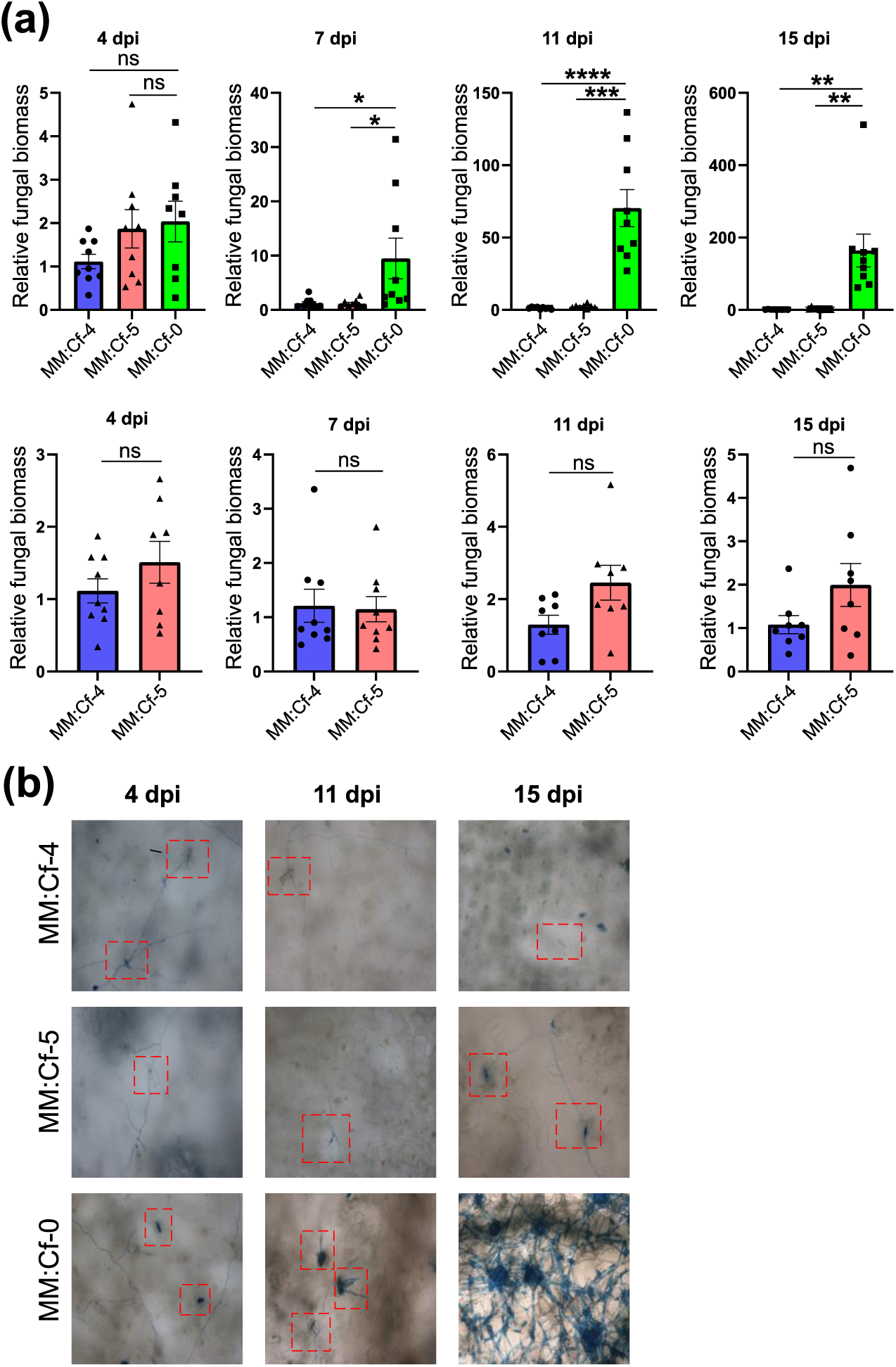
The Cf-4 and Cf-5 resistance proteins both provide robust resistance against *F. fulva*. **(a-b)** Leaves of MM:Cf-4, MM:Cf-5, and MM:Cf-0 were inoculated with *F. fulva* race 0 and leaflets were harvested at 4, 7, 11, and 15 dpi for determination of the fungal biomass **(a),** and at 4, 11, and 15 dpi, for cotton blue staining **(b). (a)** Genomic DNA was extracted from the leaves, and fungal biomass was quantified by qPCR targeting *F. fulva* actin, with normalization to tomato Rubisco. The 2^-ΔΔCT^ method was used for determining the relative biomass, with MM:Cf-4 set as a calibrator for comparison with each data point. The upper panels show a comparison of the development of the fungal biomass across all genotypes, and the lower panels show a direct comparison between the development of the fungal biomass in leaves of MM:Cf-4 and MM:Cf-5 plants only. Error bars indicate the SEM. Significant differences between the different genotypes at each time point were determined by a Student’s *t*-test (unpaired two-tailed *t*-test, p < 0.05) . * p < 0,05, ** p < 0,01, *** p < 0,001, **** p < 0,00001, ns: not significant. **(b)** Inoculated leaflets were stained with lactophenol cotton blue overnight, and cleared with ethanol. Images were taken using a Nikon Eclipse 90i microscope at 400x magnification.

Moreover, fungal development around the stomata, which are the entry points of the fungus at the lower side of the leaflets, was monitored using cotton blue staining at 4, 11 and 15 dpi for all genotypes. At 4 dpi, hyphal penetration through the stomata was observed for all genotypes. At 11 dpi, the rate of stomal colonization of the resistant genotypes was similar to that at 4 dpi, whereas conidiophore formation was observed with fungal mycelium emerging from the stomata in the susceptible MM:Cf-0 line. By 15 dpi, MM:Cf-0 showed an intense blue staining, consistent with extensive mycelial growth and abundant sporulation. However, the resistant genotypes showed no clear stomatal clogging or sporulation at any time point. Importantly, no visible differences in stomatal colonization rates were observed between MM:Cf-4 and MM:Cf-5 (Figure 7b).

Next, the expression levels of *Avr4* and *Avr5* were assessed in the resistant genotypes at 4 and 7 dpi. *Avr5* showed higher expression in both genotypes at 4 and 7 dpi, although these differences were not significant when compared with *Avr4* (Figure S11a). Because SA-related signaling contributes to resistance against biotrophic pathogens, we also quantified the expression levels of the SA-responsive gene encoding PR1 in MM:Cf-4 and MM:Cf-5, at 4 and 7 dpi using qPCR on cDNA generated from mRNA isolated from the inoculated leaves. MM:Cf-4 showed higher *PR1* transcription levels than MM:Cf-5, although this difference was not significant (Figure S11b).

## Discussion

Cf proteins are well-studied LRR-RLPs that mediate gene-for-gene resistance against *F. fulva* of tomato through recognition of matching effectors, which are secreted by *F. fulva* in the apoplastic space upon fungal ingress through the stomatal openings present at the lower side of the leaves. Cf proteins constitutively interact with the LRR-RLK SOBIR1 to compensate for the absence of a cytoplasmic kinase domain (Liebrand et al., 2013) and upon recognition of the *F. fulva* effector matching the Cf protein, the LRR-RLK BAK1 is recruited by the Cf/SOBIR1 complex (Postma et al., 2016). Subsequently, transphosphorylation between the kinase domains of SOBIR1 and BAK1 initiates, through further phosphorylation events, downstream signaling that includes the activation of RLCKs, an apoplastic ROS burst, MAPK activation, transcriptional reprogramming, and HR-related cell death, eventually resulting in resistance (van der Burgh et al., 2019; Huang et al., 2021; Huang et al., 2024; Liebrand et al., 2013; Postma et al., 2016). Cf-4, Cf-9, Cf-2, and Cf-5 are well-known RLPs (Dixon et al., 1996; Dixon et al., 1998; Jones et al., 1994; Thomas et al., 1998) and it was observed earlier that Cf-4 and Cf-9 typically induce a fast and strong cell death, whereas Cf-2 and Cf-5 trigger a slower and weaker, more chlorosis-like response (Brading et al., 2000; Cai et al., 2001; Hammond-Kosack, 1994; Rivas and Thomas, 2005).

Although the HR-related cell death is commonly associated with resistance against biotrophic pathogens (Gilchrist, 1998; Piffanelli et al., 1999), the intensity of the HR-related cell death does not determine the level of resistance in the tomato-*F. fulva* interaction, because the various Cf receptors that we study here all provide robust resistance, despite distinct cell death phenotypes (Brading et al., 2000). This observation indicates that a stronger cell death is not essential for restricting *F. fulva* growth. The molecular requirements of the Cf-4 mediated cell death and ROS burst are well defined, including the requirement of NRC2/3 and several RLCK VII-6/7/8 family members (Huang et al., 2024; Kourelis et al., 2022). However, the similarities and differences between the requirements of the individual Cf receptors, beyond their distinct HR-related cell death phenotype, have remained unknown. In this study, we selected Cf-4 and Cf-5 as a comparative model to understand receptor-specific similarities and possible differences in downstream immune responses, transcriptional reprogramming, and actual levels of resistance to *F. fulva*.

Previous studies have shown that Cf-4 requires SOBIR1, BAK1, and NRC2/3 for mounting HR-related cell death, while SOBIR1 and BAK1 are also required for Cf-4-mediated apoplastic ROS production (Huang et al., 2024; Kourelis et al., 2022; Liebrand et al., 2013; Postma et al., 2016; Zönnchen et al., 2022). However, the role of NRC2/3 in the apoplastic ROS burst has remained unknown. Here, we show that the Cf-5/Avr5-triggered cell death similarly depends on SOBIR1, BAK1, and NRC2/3, indicating that Cf-4 and Cf-5 share core signaling components to trigger HR-related cell death (Figures 1 and S1). In contrast, the apoplastic ROS burst triggered by Cf-4/Avr4 and Cf-5/Avr5 was found to require SOBIR1, but was not significantly reduced in a *bak1* or *nrc2/3* mutant background (Figures 2 and S2). This observation differs from a previous study including the Cf-4/Avr4 combination. In that study, the effect of a *bak1* knock-out was observed only when low concentrations of flg22 were applied, due to functional redundancy among the SERK family members (Zönnchen et al., 2022). Thus, the effect of the absence of BAK1 on Cf-mediated ROS production may be masked under our experimental conditions. Notably, the Cf-4/Avr4- and Cf-5/Avr5-triggered apoplastic ROS burst was found not to depend on NRC2/3, thereby supporting the existence of distinct signaling pathways for the HR-related cell death and apoplastic ROS burst. The FLS2-mediated apoplastic ROS burst has also been reported to be independent of NRC2 and NRC3 (Wu et al., 2020). Consistent with this observation, RLCK VII-6 subfamily members are required for a full apoplastic ROS burst, but are not essential for the HR-related cell death (Huang et al., 2024). Moreover, the Cf-9/Avr9 combination also triggers a biphasic apoplastic ROS burst, similar to that observed for Cf-4/Avr4 and Cf-5/Avr5, indicating that the different Cf proteins trigger a biphasic ROS burst in *N. benthamiana*. Despite clear differences in the HR-related cell death phenotypes among Cf receptors, we did not detect significant differences in the magnitude of the apoplastic ROS burst that they trigger (Figure S3). Together, these results demonstrate that the variation in the strength of the cell death triggered by the different Cf receptors, is uncoupled from apoplastic ROS production in *N. benthamiana*.

In tomato, Avr4 triggers a strong cell death in MM:Cf-4 plants, whereas Avr5 induces a noticeably weaker cell death in MM:Cf-5. Neither effector triggered an HR-related cell death in plants carrying non-matching receptors or in MM:Cf-0, confirming the specificity of the responses (Figure 3a). Next, MAPK activation dynamics were assessed at different time points in tomato. Both Cf-4/Avr4 and Cf-5/Avr5 induced an early MAPK activation within 15 min after challenge, with no significant differences in the intensity (Figures 3b and S4a). However, a clear difference emerged at later stages, when Cf-5/Avr5 elicited a more sustained and prolonged MAPK activation when compared with Cf-4/Avr4. In contrast, infiltration of a solution of Avr4 or Avr5 triggered only weak and transient MAPK activation at 15 mins in leaves of MM:Cf-0 plants (Figure 3b), whereas flg22 induced a strong MAPK response at 15 mins, which rapidly decreased at later time points (Figure S4b).

It has been shown that flooding of the apoplast with water leads to transcriptional changes and phosphorylation of calcium-dependent kinases (CDPKs) (Witte et al., 2010). In addition, mechanical damage caused by the infiltration procedure itself might activate MAPKs (Hõrak, 2020). Therefore, the weak and transient MAPK activation in MM:Cf-0 plants might reflect a stress-related activation. Together, these results indicate that the dynamics of MAPK activation differ not only between Cf-4 and Cf-5, but also between Cf-mediated and FLS2-mediated responses in tomato, with both Cf proteins displaying MAPK activation profiles that are distinct from those induced by flg22. Similar differences in MAPK activation kinetics have been reported in *A. thaliana*, where the AvrRpt2 effector of the pathogenic bacterium *Pseudomonas syringae*, which is indirectly perceived by the cytoplasmic resistance protein RESISTANCE TO *PSEUDOMONAS SYRINGAE* (RPS2) through the cleavage of RPM1-INTERACTING PROTEIN (RIN4) (Axtell and Staskawicz, 2003), triggers a more sustained MAPK activation than flg22. Moreover, the same study showed distinct MAPK activation dynamics between AvrRpt2 and AvrRps4 (Tsuda et al., 2013), indicating that different receptor–effector combinations might have different MAPK activation dynamics. Furthermore, it has been shown that different effectors differentially regulate MAPK activation in alfalfa (Cardinale et al., 2000). A previous study reported prolonged MAPK activation by Cf-4/Avr4 in *N. benthamiana* (Huang et al., 2022). However, such prolonged activation was not observed in tomato in this study. In *N. benthamiana*, MAPK activation remained strong even at 90 minutes, whereas in tomato, the signal declined much earlier. Therefore, it is likely that species-specific differences occur in the MAPK activation dynamics by the Cf-4/Avr4 combination.

Next, we compared the intensity and timing of the apoplastic ROS burst triggered by Cf-4 and Cf-5 in tomato, upon their elicitation. To validate the apoplastic ROS assay in tomato, we first tested the method in tomato using flg22, which resulted in a clear ROS burst when compared with the MQ negative control (Figure S5a). The same assay was then applied to tomato, where flg22 induced a strong ROS burst in leaf discs taken from the wild-type Rio Grande cultivar, but not when such discs were taken from the *rbohb* mutant (Figure S5b), thereby confirming that the method reliably measures the RBOHB-dependent apoplastic ROS production in tomato leaves. In MM:Cf-4 and MM:Cf-5 tomato, respectively both Avr4 and Avr5 induced a ROS production that was clearly distinguishable from the MQ treatment (Figure 4a). However, no significant differences in the ROS burst intensity and timing were detected when the Cf-4/Avr4 and Cf-5/Avr5 combinations were compared (Figure S6). Although Avr9 triggered a clear biphasic ROS burst in *N. benthamiana* transiently expressing Cf-9 (Figure S3), it was difficult to distinguish the Avr9-induced ROS from the MQ controls in MM:Cf-9 and *Hcr9-9D*-stable transgenic tomato plants (Figure S7). Moreover, higher concentrations of Avr4 and Avr5 were required to induce clear ROS bursts in tomato, when compared to the concentrations used in *N. benthamiana*. Together, these results suggest that effector-triggered apoplastic ROS responses are comparatively weaker in tomato.

Unexpectedly, Avr4 and Avr5 also trigger an apoplastic ROS burst in the negative controls, including MM:Cf-0 and tomato plants carrying a non-matching receptor, without any significant difference from tomato plants carrying the matching receptor (Figures 4b and S8). Furthermore, both Avr4 and Avr5 effectors induce ROS production in Rio Grande wild-type plants, but not in the *rbohb* mutant (Figure S9), indicating that these ROS bursts are RBOHB-dependent. Moreover, the Cf-4/Avr4 triggered apoplastic ROS burst in *N. benthamiana* is dependent on RBOHD (Landeo Villanueva, 2023). Both Avr4 and Avr5 were produced in the yeast *P. pastoris*, purified using a His-tag affinity column, and subsequently dialyzed using a 3.5 kDa molecular weight cut-off membrane. Although this purification strategy substantially reduces the presence of co-purifying yeast-derived molecules, the possible contribution of residual *P. pastoris*–derived components to the observed apoplastic ROS burst was still considered, particularly given that *P. pastoris* secretes a relatively small number of proteins during methanol induction (Burgard et al., 2020). Furthermore, classical PAMPs potentially originating from *P. pastoris*, such as chitin, are known to trigger a rapid apoplastic ROS burst, for example, within minutes of treatment in tomato (Li et al., 2024; Rahman et al., 2020). Similar rapid ROS kinetics have been reported for β-glucans, both of which trigger ROS with kinetics similar to flg22 (Wanke et al., 2020). In contrast, the apoplastic ROS responses induced by Avr4 and Avr5 are delayed when compared to the flg22-triggered ROS burst (Figures 4 and S5). Consistent with these observations, our MAPK activation assays in MM:Cf-0 revealed only a weak and transient MAPK phosphorylation at 15 min following treatment with Avr4 or Avr5 (Figure 3c), in contrast to the strong, early and transient MAPK activation induced by flg22 in these plants (Figure S4b). Together, we propose that the apoplastic ROS burst that is triggered independently of a matching Cf protein might not be related to PAMPs originating from *P. pastoris* but could have something to do with the virulence function of the effectors. For example, previous studies have shown that Avr9 binds with high affinity to the plasma membrane of tomato leaf cells even in the absence of Cf-9, and suggested the presence of additional receptor-like kinases or guarded virulence targets, like a high-affinity binding site (HABS) involved in Avr9 binding (Kooman-Gersmann et al., 1996; Zhao et al., 2022). Our results suggest that Avr4 and Avr5 may similarly trigger an apoplastic ROS through Cf-independent perception mechanisms in tomato. However, previous studies have shown that silencing of *Cf-4* in *N. benthamiana:Cf-4* reduces the apoplastic ROS burst (Landeo Villanueva, 2023), indicating that the receptor-independent apoplastic ROS burst is specific for tomato. Further experiments using proteins purified from *P. pastoris* carrying an empty vector will help to clarify these observations. Note that the Avr4 and Avr5 protein are carrying a His tag for affinity column purification. Removing the His tag after their purification might be worthwhile to ensure that recognition of this tag by the plant is not triggering the ROS burst that is observed in the absence of the matching Cf proteins.

Transcriptomic analysis of the Cf-4/Avr4- and Cf-5/Avr5-triggered responses of tomato leaves showed that both immune pathways involve a largely overlapping core immune transcriptional program, but that the Cf-5/Avr5 combination consistently induces broader and stronger transcriptional reprogramming than the Cf-4/Avr4 combination at the tested time points. Furthermore, both combinations also resulted in quantitative and qualitative differences in downstream immune signaling events (Figure 5 and S10). For example, the differential regulation of phytohormone-related genes between the Cf-4/Avr4 and Cf-5/Avr5 combinations might reflect differences in the timing of hormone signaling (Figure 6a). SA signaling is known to be induced by Cf proteins during elicitor treatment or upon pathogen infection (Etalo et al., 2013; van Kan, 2012; Rivas and Thomas, 2005), although SA-related immune signaling is not required for resistance against *F. fulva* (Brading et al., 2000). Cf-4/Avr4 may trigger an earlier, or a more rapid SA accumulation already before the 3h sampling point. Consequently, transcriptional regulation of SA-associated genes may no longer be, or may not be strongly detected anymore at 3h in the Cf-4/Avr4 combination (Figure 6a). Consistent with this idea, JA signaling appeared to be more tightly regulated in the Cf-4/Avr4 combination, as evidenced by a stronger transcriptional upregulation of JA suppressors, such as the *JAZ* and *CYP94* genes (Chini et al., 2007; Koo et al., 2011; Yang et al., 2024) (Figure 6a). SA signaling might be induced at early time points, followed by activation of JA signaling, which is explained by the well-known SA–JA antagonism. This JA response might subsequently again be reduced by the up-regulation of JA suppressors.

In contrast, the Cf-5/Avr5-triggered immune response is associated with a broader and more sustained transcriptional regulation across the SA-, ET-, and JA-regulated pathways (Figure 6a). Genes encoding pathogenesis-related (PR) proteins were clearly upregulated in both immune pathways, in most cases largely overlapping but differing quantitatively, with generally stronger induction in the Cf-5/Avr5 combination. Notably, the gene encoding Solyc02g077370, which is PTI5, has been reported to be induced during pathogen attack (Du et al., 2015; Meller Harel et al., 2014; Vega et al., 2015), and was specifically upregulated in the Cf-5/Avr5 combination, further supporting differences in downstream defense signaling between the two pathways (Figure 6b). Transcriptional upregulation of genes encoding various components of the MAPK cascades was largely shared between Cf-4/Avr4 and Cf-5/Avr5. However, the Cf-5/Avr5 combination resulted in a more sustained transcriptional activation of several of these components-encoding genes, indicating a more prolonged and sustained initiation of signal transduction (Figure 6).

*WRKYs* were similarly regulated in both pathways, but each Cf/Avr combination also resulted in the activation of a distinct subset of *WRKYs*, and with many shared *WRKYs* showing a stronger induction in the Cf-5/Avr5 combination (Figure S10a). This observation suggests that an enhanced transcriptional amplification takes place in Cf-5/Avr5-triggered immunity. The transcriptional regulation of genes associated with the apoplastic ROS burst also showed a substantial overlap between the two combinations, but again with a stronger induction of the genes in the Cf-5/Avr5 combination (Figure 6d). Different RBOH-encoding genes are upregulated during immune signaling activation. In tomato, several *RBOHs* are induced during *Botrytis cinerea* infection, although only *RBOHB* is required for resistance against this necrotrophic fungus (Li et al., 2015). Moreover, in *A. thaliana*, *RBOHD* and *RBOHF* displayed differential expression patterns during infection by *B. cinerea* (Morales et al., 2016), suggesting that the different RBOH-encoding genes might have distinct roles in immune signaling. Notably, *RBOHA* was specifically transcriptionally upregulated in Cf-5/Avr5. *RBOHA* is, for example, upregulated during *B. cinerea* infection (Li et al., 2015), indicating that Cf-5/Avr5 may engage in additional ROS-related defense mechanisms, thereby providing yet another piece of evidence of a difference in immune signaling downstream of Cf-4/Avr4 and Cf-5/Avr5.

Overall, Cf-5/Avr5-triggered immunity exhibited a stronger and broader transcriptional response when compared to Cf-4/Avr4, which is consistent with our MAPK activation results. However, these differences may also be influenced by the distinct HR-related cell death phenotypes associated with the two immune pathways. Cf-4/Avr4 induces a strong and fast cell death response, whereas the Cf-5/Avr5 combination triggers a relatively slow chlorosis. Extensive and fast cell collapse as a result of Cf-4 challenge by Avr4 may limit the possibilities for a sustained transcriptional reprogramming, while the absence of a fast cell death response in Cf-5/Avr5 could allow a longer temporal window for a massive and sustained transcriptional reprogramming. Consistent with this idea, our analysis of the transcriptional regulation of PCD-related genes revealed that genes encoding BAX1 inhibitor (BI) proteins, which function as suppressors of PCD (Ishikawa et al., 2011; Watanabe and Lam, 2009), were continuously upregulated in Cf-5/Avr5 (Figure S10b). This sustained upregulation suggests that Cf-5/Avr5 may actively promote pro-survival mechanisms that limit the strong cell death phenotype, thereby maintaining cellular integrity and enabling prolonged and robust immune signaling. Together, these observations raise the possibility that differential regulation of cell survival and cell death contributes to the distinct transcriptional outputs and macroscopic phenotypes observed between Cf-4/Avr4 and Cf-5/Avr5-triggered immunity. Overall, both Cf-4 and Cf-5 provide full resistance against *F. fulva,* indicating that both immune strategies are effective in preventing pathogen infection (Figure 7). Many studies have shown that *F. fulva* can overcome Cf-mediated resistance through deletion or mutation of effector genes (Stergiopoulos et al., 2007). This raises the question of whether differences in immune response dynamics contribute to the long-term durability of individual *Cf* genes.

## Materials and Methods

### Plant growth conditions

*N. benthamiana* wild-type (WT) and the *sobir1/sobir1-like* (Huang et al., 2021)*, bak1* (Zönnchen et al., 2022)*, nrc2/3_3.3.1* (Kourelis et al., 2022)*, eds1* (Zönnchen et al., 2022) and *pad4* (Zönnchen et al., 2022) *N. benthamiana* knock-out plants were grown in a climate chamber under a 16-hour light cycle at 25°C, and 8-hour dark cycle at 21°C, and at 70% humidity. MM:Cf-0, MM:Cf-4, MM:Cf-5, Rio Grande and Rio Grande *rbohb* (Leuschen-Kohl et al., 2025) tomato were grown in a controlled climate chamber at 24°C and 75% relative humidity (RH), with a photoperiod of 16 hours and 8 hours of darkness.

### Binary Vectors for Agrobacterium-mediated transformation

Binary vectors for transient expression of Cf-4 (Liebrand et al., 2013), Avr4 (van der Hoorn et al., 2000), Cf-5, and Avr5 (Kourelis et al., 2020) were described previously. Cf-4 and codon-optimized (CO) Cf-5 were expressed from the same binary vectors. For this, Cf-5:CO was amplified from the provided template plasmid using Phusion Hot Start II High-Fidelity Polymerase (Promega), using the primers indicated in Table S1, and ligated into pENTR™/D-TOPO^®^. Subsequently, the coding sequences of all constructs were transferred from this vector to the pBIN-35S binary vector (Sol 2095) using a Gateway™ LR Clonase™ II Enzyme mix (Thermo Fisher Scientific). The binary vector provides a C-terminal eGFP tag, and the constructs are referred to as “pBIN-35S::”.

### Production of the Avr5 protein in the yeast *Pichia pastoris*

Recombinant His-tagged Avr5 protein production was performed in the yeast *Pichia pastoris* (van den Burg et al., 2001). The culture supernatant was concentrated using a VIVAFLOW 200 device, with a 5000D molecular weight cut-off (MWCO) membrane (Sartorius), at 4°C. The concentrated supernatant was then dialyzed against equilibration buffer (25mM Tris, 25mM NaCl, 10mM imidazole, pH=7.5), using Spectrum™ Labs Spectra/Por™ 7 pre-treated dialysis tubing (Spectrum Labs; MWCO 3.5 kDa). The Avr5 protein was subsequently purified using a Biologic LP low pressure chromatography system (BIO-RAD), with a column containing His60 nickel beads Ni-NTA resin for purification of His-tagged proteins (Takara Bio). A 10 mM to 200 mM gradient of imidazole in elution buffer (25 mM Tris, 25 mM NaCl, pH=7.5), over a period of 20 minutes at a flow of 1 ml/min, was used to elute the Avr5 protein from the column, after which the solution of purified Avr5 protein was dialyzed against MQ. Determination of the Avr5 protein concentration was performed by the Quick Start Bradford Protein Assay (BIO-RAD), and the presence of the protein was verified by immunoblotting using 6x-His Tag Monoclonal Antibody (Invitrogen, MA1-21315).

### HR-related cell death assay

HR-related cell death assays in different *N. benthamiana* lines were carried out using the previously described Agrobacterium-mediated transformation assay (van der Hoorn et al., 2000). Shortly, Cf-4 and Cf-5, all tagged with eGFP, were agroinfiltrated in leaves of the different *N. benthamiana* lines, in combination with their matching effectors, Avr4 and Avr5 (OD_600_=0.8), respectively. The agroinfiltrated leaves were imaged at 7 dpi using the red-light imaging method with a Chemidoc XRS system (BIO-RAD) and ImageLab was used to quantify the intensity of the HR-related cell death (Landeo Villanueva et al., 2021). HR-related cell death assays in tomato MM:Cf-0, MM:Cf-4, and MM:Cf-5 were performed using solutions of the pure Avr4 and Avr5 effectors. For this, solutions of 5 µM Avr4 and Avr5 were infiltrated in leaves of 5-week-old tomato plants, while MQ and their non-matching effectors were used as negative controls. At 7 dpi, leaves were imaged, and the intensity of the HR-related cell death was quantified as described above

### Apoplastic ROS burst assay

The apoplastic ROS burst in tomato was assessed using a luminol-based assay. For this, 5 mm leaf discs were taken from leaflets of the first or secondary fully expanded leaf of 4-or 5-week-old tomato plants with a biopsy punch and placed in a 96-well white micro-titer plate (Greiner Bio-One), containing 50 µL of MQ. The leaf discs were incubated overnight in the dark at room temperature, after which the MQ was replaced by fresh MQ (50 µL), in which the leaf discs were incubated for at least one hour in the dark at room temperature.

Then, 50 µl of the reaction mix, containing 50 µM luminol L-012 (FUJIFILM), 10 μM horseradish peroxidase, Avr4, Avr5, Avr9, and flg22, all at a concentration of 0.1 μM for *N. benthamiana* and 1 μM for tomato, were added to the wells. MQ was used as a negative control. The apoplastic ROS burst was recorded over a period of 5 hours, in reading cycles of 2 minutes each, using a CLARIOstar plate reader (BMG Labtech).

### MAPK activation assays

To assess possible differences in MAPK activation patterns between Cf-4 and Cf-5, 5 μM solutions of Avr4 and Avr5 were infiltrated into leaves of MM:Cf-4 and MM:Cf-5 tomato plants, respectively. The effectors were also infiltrated in MM:Cf-0 as a negative control. For the flg22 peptide the concentration was 1 μM. After infiltration, 10 leaf discs per treatment were harvested at 0, 15, 30, 60, and 90 mins, and immediately stored in liquid nitrogen. The leaf discs were powdered using a laboratory mixer (Retsch GMBH, Haan, Germany) for 3 minutes at 30 revolutions/s, after which the total proteins were extracted using an extraction buffer containing 150 mM NaCl, 1% IGEPAL^®^ CA-630, 50 mM Tris-HCl (pH 8.0), and 1 protease inhibitor cocktail tab per 50 ml. Proteins were separated by SDS-PAGE and subsequently subjected to immunoblotting. p44/p42 MAPK (Erk1/2) antibodies (NEB), followed by an anti-Rabbit-HRP secondary antibody, were used to detect activated MAPKs.

### Inoculation assays

*F. fulva* race 0 [CBS131901, (de Wit et al., 2012)], secreting both Avr4 and Avr5, was grown on potato-dextrose agar (PDA), with 1.5% technical agar extra, at room temperature for 4 weeks. Conidia were harvested from the plates in tap water, and their final concentration was adjusted to 1x10^6^ conidia/ml. *F. fulva* inoculations were performed as previously described (Ökmen et al., 2013). Inoculated leaves were harvested 4, 7, 11, and 15 dpi and subjected to the determination of fungal biomass and relative gene expression levels by qPCR.

### Determination of fungal biomass and gene expression levels by qPCR

Fungal biomass was quantified by qPCR on genomic DNA (gDNA), whereas *Avr4*, *Avr5*, and *PR1a* expression levels were assessed by qPCR on cDNA. The genomic DNA was extracted from *F. fulva*-inoculated leaves using a previously described method (Murray and Thompson, 1980) and total RNA was isolated using the Maxwell^®^ RSC Plant RNA kit (Promega). cDNA was synthesized from total RNA using M-MLV Reverse Transcriptase (Promega) and an oligo(dT) primer. For the qPCR, the protocol of the SensiMix™ SYBR® No-ROX Kit (Meridian Bioscience) was followed, including the appropriate primers (Table S1).

### Cotton blue staining

Fungal colonization of the inoculated leaves was assessed using cotton blue staining. Inoculated leaves were harvested at 4, 11, and 15 dpi, and incubated in a lactophenol blue solution (Sigma Aldrich), with gentle agitation overnight. After that, the leaves were destained by washing in ethanol for 24-48 h. Random sections of the stained leaves were imaged using a Nikon Eclipse 90i microscope at 400x magnification.

### Preparation of leaf samples for full transcriptome analysis by RNA-seq

Leaves of MM:Cf-4 and MM:Cf-5 tomato plants were infiltrated with a solution of 0,5 µM Avr4 and Avr5, respectively, after which the samples were harvested at 0, 3 and 7h, and directly stored in liquid nitrogen. Total RNA extraction, library preparation, sequencing and primary RNA-seq data analysis were performed by the company Biomarker Technologies (BMK) GmbH, Münster, Germany.

## Acknowledgements

Henriek Beenen provided the pPIC Avr5 plasmid for Avr5 protein production in yeast. Sergio Landeo Villanueva and Ciska Braam contributed to the production of the Avr4 and Avr5 proteins. Christiaan Schol helped with the *F. fulva* inoculation assay. Prof. Sophien Kamoun provided the *nrc2/3_3.3.1* knock-out mutant of *N. benthamiana,* Prof. Greg Martin provided the tomato Rio Grande and Rio Grande *rbohb* knock-out lines and Johannes Stuttmann provided the *bak1* knock-out mutant of *N. benthamiana*. We thank Unifarm personnel for taking care of the plants and we thank the SOL-team for fruitful discussions. E.B. is supported by The Ministry of National Education of the Republic of Türkiye.

## Author Contributions

E.B. and M.H.A.J.J. designed the work and wrote the manuscript. E.B. and H.A.C performed experiments. All authors read and approved the manuscript.

