## Supplemental Data 1 for "Cf-4- and Cf-5-Triggered Plant Immunity: Similarities and Differences"

**Supplementary Data**

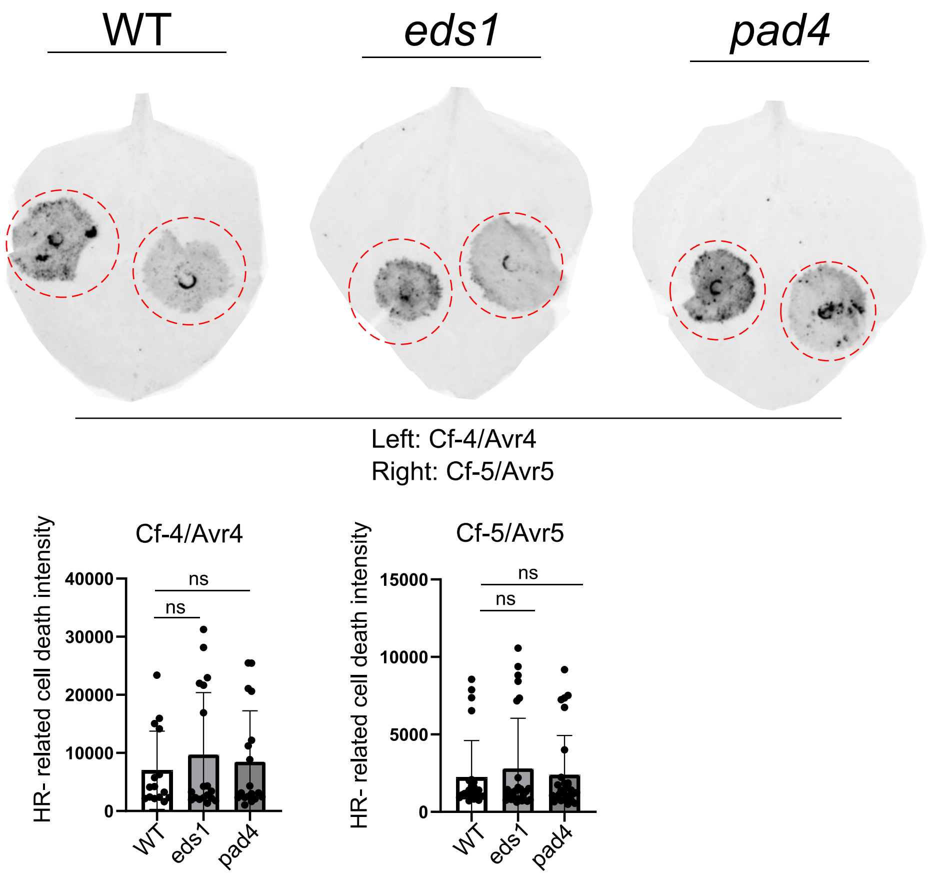


**Figure S1: Both EDS1 and PAD4 are not essential for the Cf-4/Avr4- and Cf-5/Avr5-triggered HR-related cell death.** Cf-4/Avr4 and Cf-5/Avr5 were transiently co-expressed by agroinfiltration in leaves of the indicated *N. benthamiana* lines ($OD_{600}$=0.8), after which the infiltrated leaves were imaged using red-light imaging at 6 dpi. ImageLab was used for quantification of the intensity of HR-related cell death, and an ANOVA/Dunnett's multiple comparison test was used to assess significant differences. The experiment was repeated three times; ns, not significant. WT, wild-type.

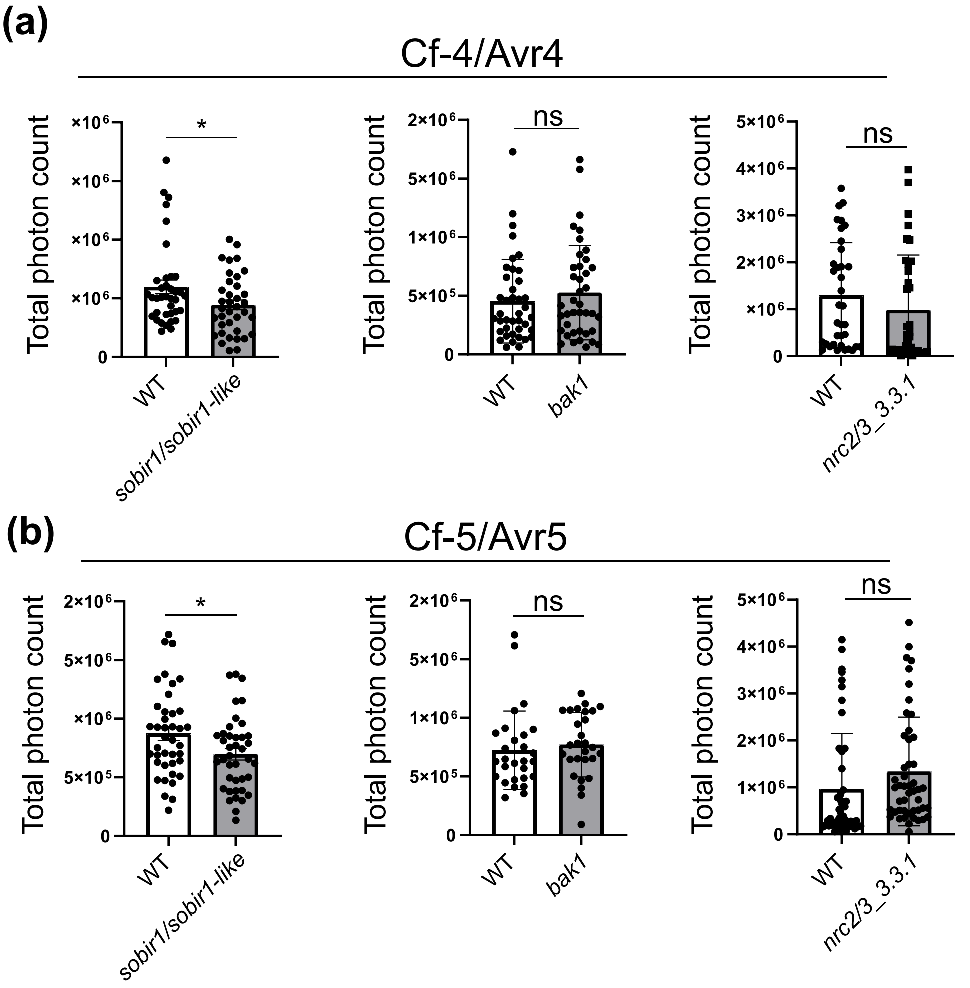


**Figure S2: Both NRC2 and NRC3 are not required for the Cf-4/Avr4- and Cf-5/Avr5-triggered ROS burst. (a)** Cf-4 and **(b)** Cf-5 were transiently expressed in leaves of the indicated lines of *N. benthamiana* ($OD_{600}$=0.1), and at 2 dpi, leaf discs were challenged by incubation with Avr4 **(a)** and Avr5 **(b)**, both at a concentration of 0.1 µM, over a total period of 5 hours. A luminol-based assay was used to measure the apoplastic ROS burst. The experiment was repeated at least three times. To assess the intensity of the apoplastic ROS burst, the total photon counts (the area under the curves) were calculated from the sum of three independent experiments. A Student's *t*-test was used to assess significant differences between the wild-type (WT) and mutant lines (unpaired two-tailed *t*-test, $P < 0.05$). Error bars represent the SD. * $p < 0,05$; ns, not significant.

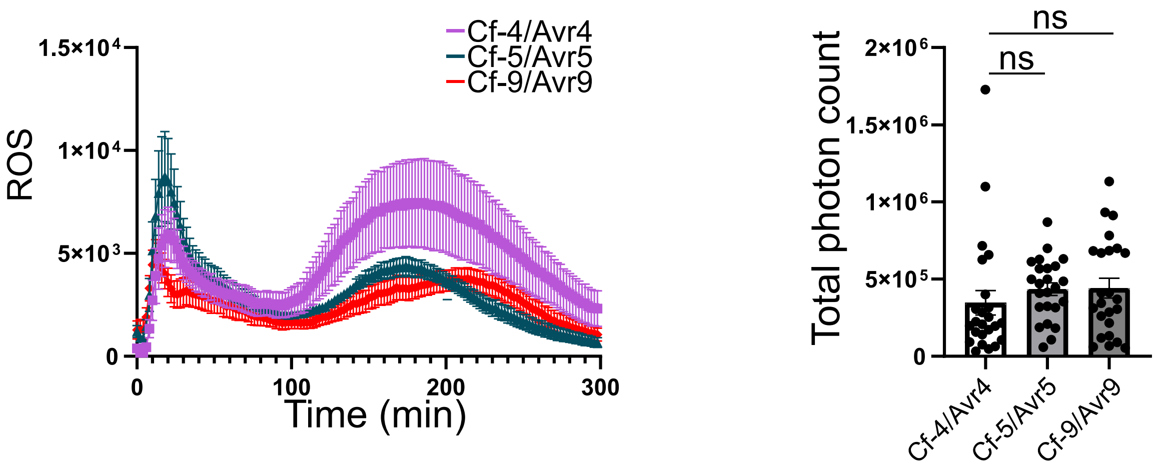


**Figure S3: Cf-4, Cf-5, and Cf-9 trigger a biphasic ROS burst with comparable overall intensity upon challenge with their matching effector, in leaves of *N. benthamiana*.** Leaf discs of *N. benthamiana* plants transiently expressing Cf-4, Cf-5, and Cf-9 were incubated with a 0.1 µM solution of Avr4, Avr5, and Avr9, respectively, for 5 hours. The apoplastic ROS burst was measured using a luminol-based assay (left) and the total photon counts (the area under the curves), were calculated to quantify the intensity of the ROS burst (right). An ANOVA/Dunnett's multiple comparison test was used to assess significant differences. Error bars show the SEM; ns, not significant.

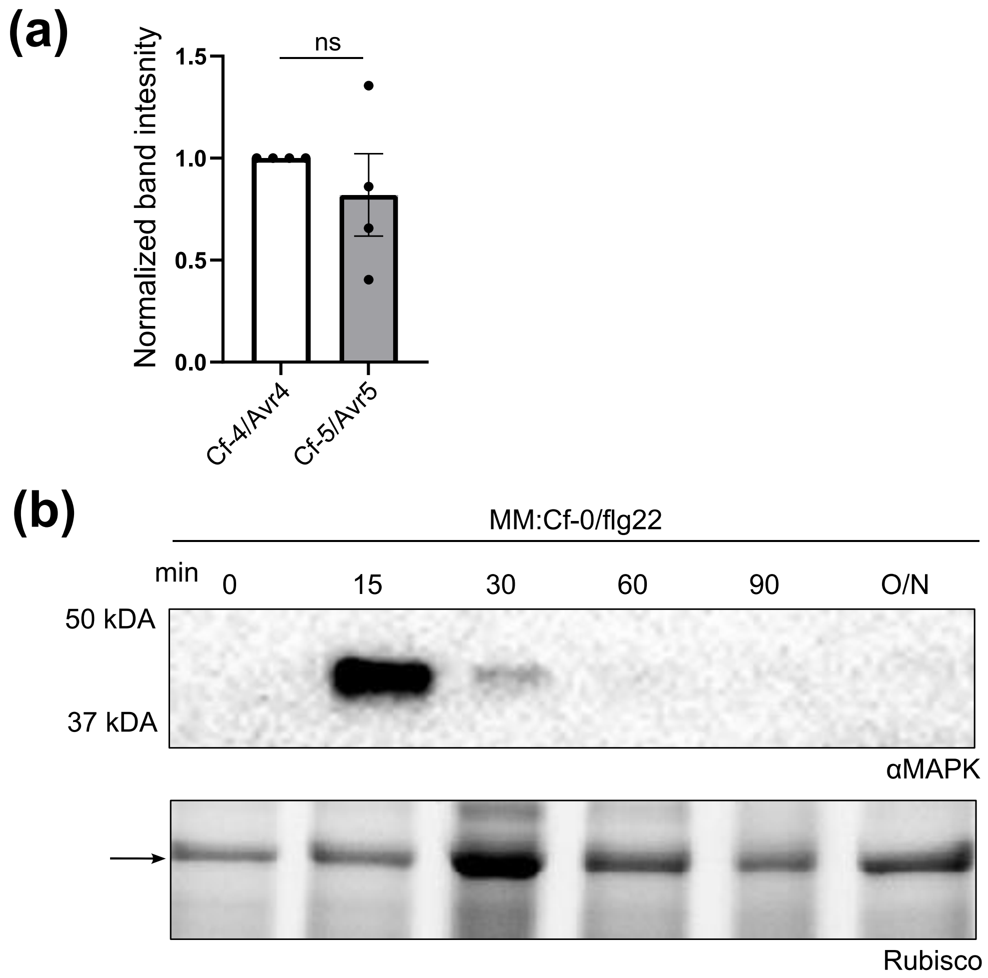


**Figure S4: Cf/Avr- and FLS2/flg22-triggered MAPK activation in tomato leaves. (a)** The intensity of the bands representing MAPK activation by Cf-4/Avr4 and Cf-5/Avr5 at 15 min after challenge, as shown in Figure 3, which were detected using antibodies against phosphorylated MAPKs, was normalized to the intensity of the Rubisco bands. **(b)** A solution of 1 μM flg22 was infiltrated into leaves of MM:Cf-0 tomato plants, after which leaves were sampled at 0, 15, 30, 60, 90 mins and 24 hours (O/N). Total proteins were extracted and subjected to immunoblotting, after which antibodies against phosphorylated MAPKs were used to detect the activated MAPKs. Total protein loading is indicated by the intensity of the Rubisco band. The experiment was repeated three times, and a representative result is shown.

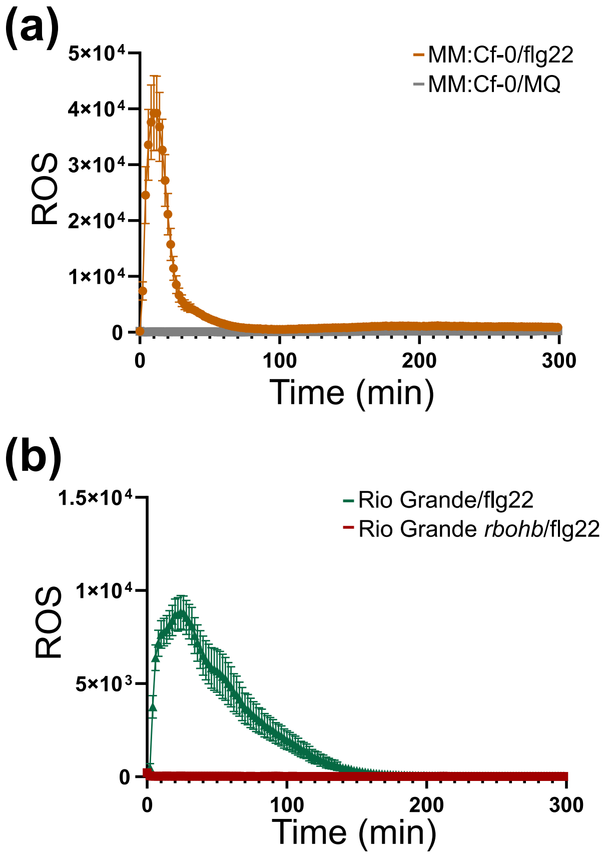


**Figure S5: FLS2/flg22 triggers a monophasic ROS burst in leaves of MM:Cf-0 and tomato Rio Grande (RG), but not in an RG *rbohb* knock-out line.** Leaf discs from MM:Cf-0 **(a),** RG and RG *rbohb* **(b)** were challenged with a 1 μM solution of flg22 over a period of 5 hours and ROS was detected using a luminol-based assay. MQ served as a negative control for MM:Cf-0. The experiment was repeated three times, and representative results are shown. The error bars represent the SEM.

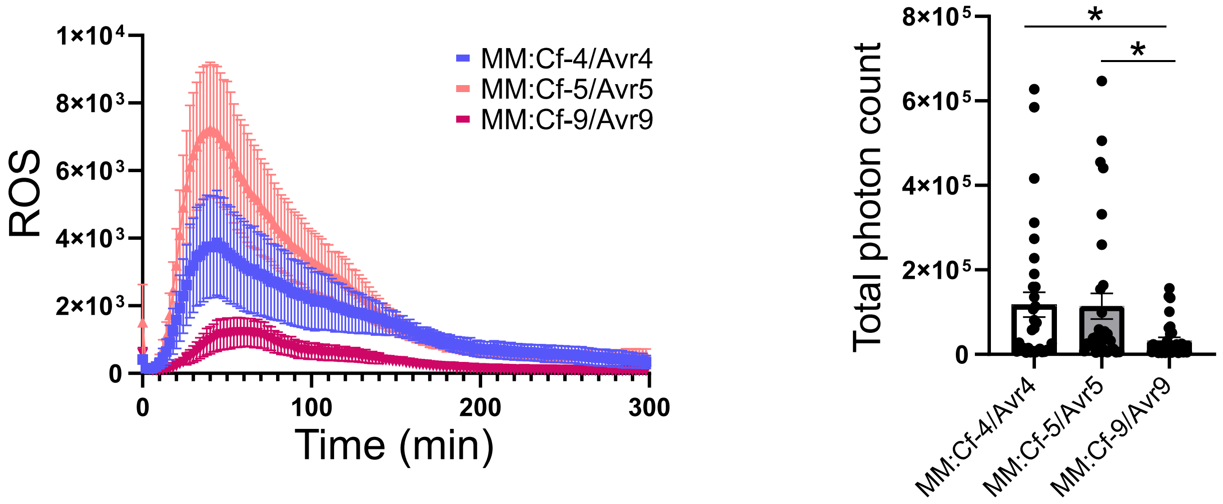


**Figure S6: The Cf-9/Avr9 combination results in a less intense apoplastic ROS burst than Cf-4/Avr4 and Cf-5/Avr5, in leaves of tomato.** Leaf discs from MM:Cf-4, MM:Cf-5, and MM:Cf-9 were challenged with a solution of 1 μM Avr4, Avr5, and Avr9, respectively, over a period of 5 hours and ROS was detected using a luminol-based assay. The experiment was repeated three times, and representative images are shown for the obtained ROS profiles (left). Total photon counts (the area under the curves) were calculated to quantify the overall ROS burst intensity (right). ANOVA/Dunnett's multiple comparison was used to assess significant differences and the error bars show the SEM; * $p < 0,05$.

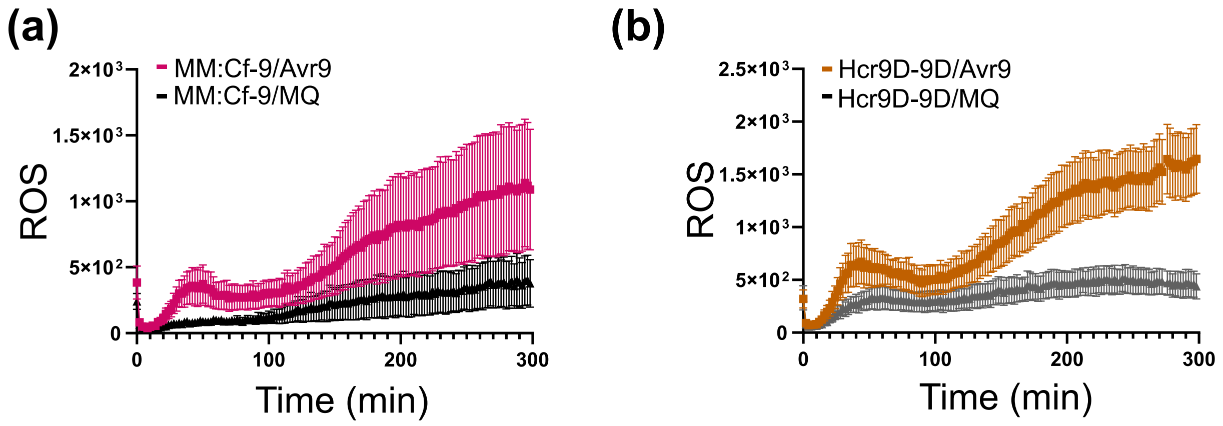


**Figure S7: The Avr9-triggered apoplastic ROS burst in MM:Cf-9 and Hcr9-9D tomato shows a less clear distinction from the MQ treatment.** Leaf discs were taken from MM:Cf-9 **(a)** and Hcr9-9D tomato plants **(b)**, and they were challenged with a solution of 1 μM Avr9 over a period of 5 hours. ROS was detected using a luminol-based assay, in which MQ was used as a negative control. Error bars show the SEM.

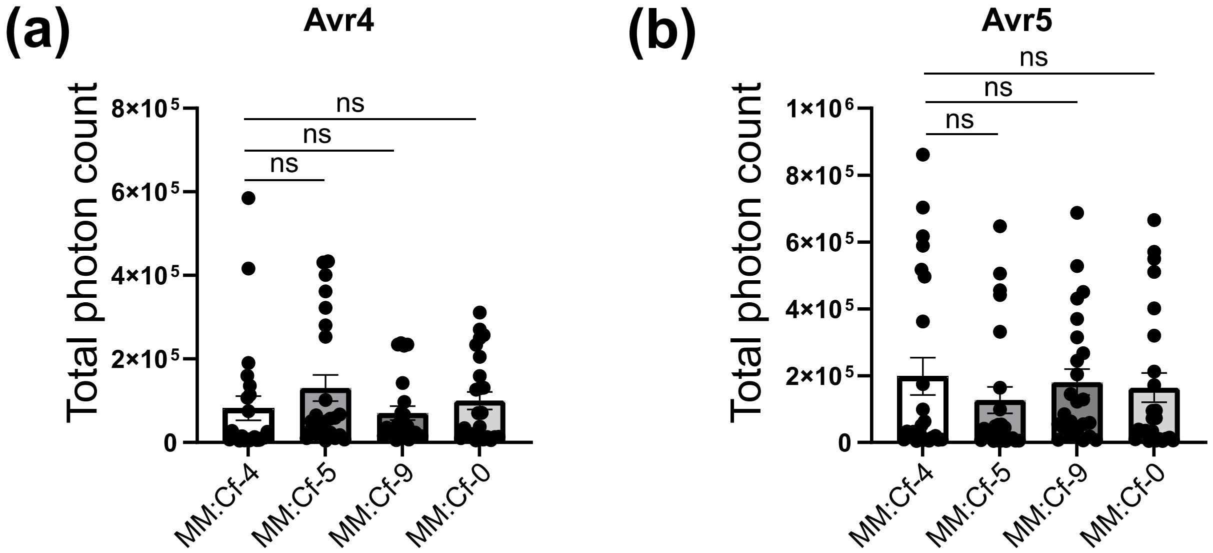


**Figure S8: Avr4 and Avr5 trigger an apoplastic ROS burst also in tomato lines lacking the matching *Cf* gene, without significant differences.** Leaf discs were taken from MM:Cf-4, MM:Cf-5, MM:Cf-9, and MM:Cf-0 tomato plants and they were challenged with a 1 µM solution of Avr4 **(a)** and Avr5 **(b)** over a period of 5 hours, during which ROS was detected using a luminol-based assay. Total photon counts (the area under the curves) were calculated to quantify the intensity of the ROS burst. An ANOVA/Dunnett's multiple comparison test was used to assess significant differences. Error bars show the SEM, ns: not significant.

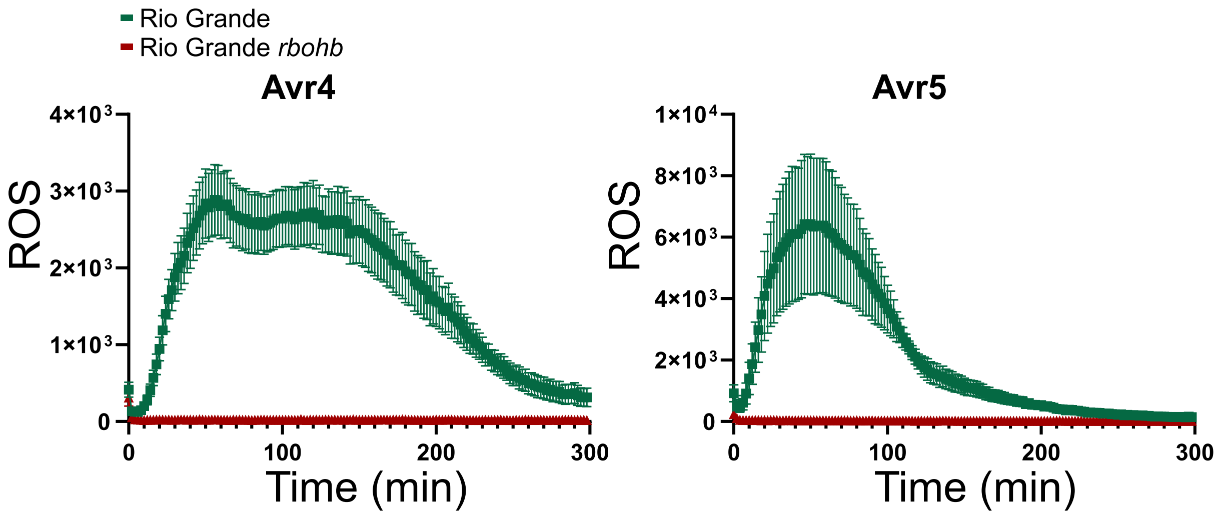


**Figure S9: The Avr4- and Avr5-triggered ROS burst is fully RBOHB-dependent in tomato Rio Grande.** Leaf discs, taken from tomato Rio Grande (RG) and a RG *rbohb* knock-out, were challenged with a solution of 1 µM Avr4 (left) and Avr5 (right) for 5 hours, and a luminol-based assay was used to measure the apoplastic ROS burst. The experiment was repeated three times, and representative images are shown. Error bars represent the SEM.

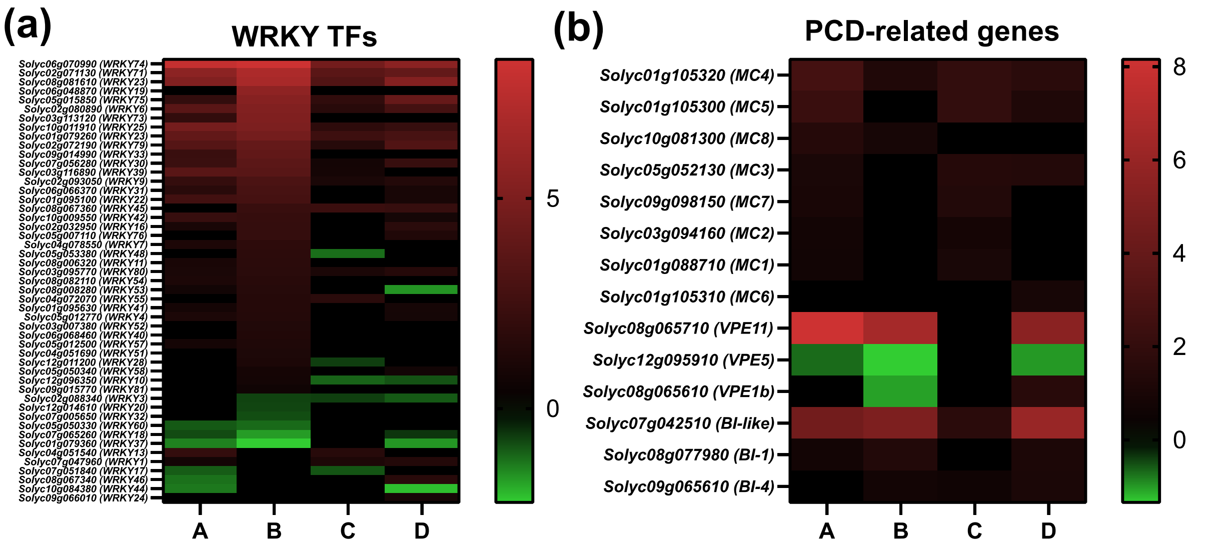


**Figure S10: The Cf-4/Avr4- and Cf-5/Avr5-triggered immune responses share a large proportion of transcriptionally differentially regulated *WRKYs* and selected PCD-related genes.** The transcription levels of genes encoding **(a)** WRKY transcription factors (TFs) and **(b)** selected PCD-related genes were visualized using heatmaps to assess qualitative and quantitative differences in changes in gene expression between Cf-4/Avr4 and Cf-5/Avr5. A, Cf-4/Avr4 at 3h; B, Cf-5/Avr5 at 3h; C, Cf-4/Avr4 at 7h; D, Cf-5/Avr5 at 7h.

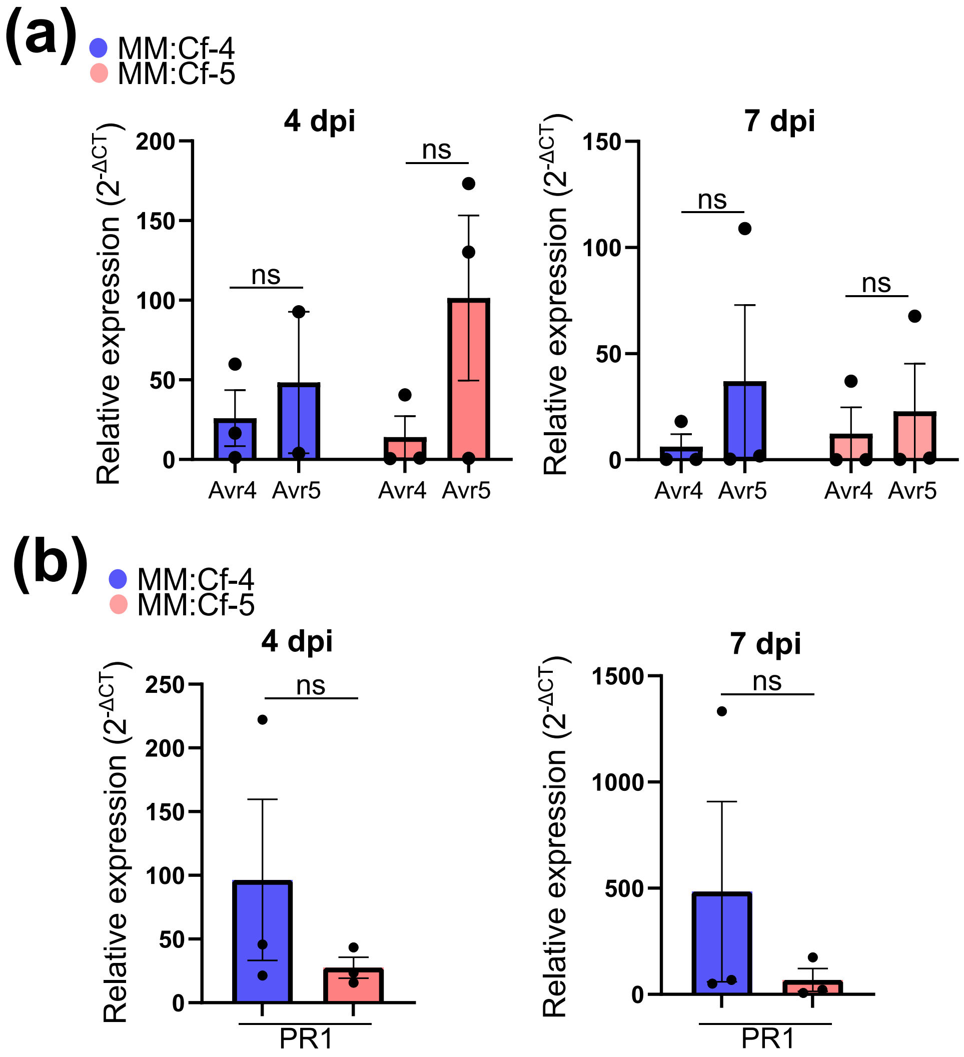


**Figure S11: *Avr4*, *Avr5*, and *PR1* are similarly expressed in tomato MM:Cf-4 and MM:Cf-5 upon inoculation with a race 0 of *F. fulva*. (a-b)** MM:Cf-4 and MM:Cf-5 tomato plants were inoculated with *F. fulva* race 0, secreting both Avr4 and Avr5. The inoculated leaves were harvested at 4 and 7 dpi, followed by RNA extraction, and cDNA synthesis. Relative expression levels of *Avr4* and *Avr5* **(a)** and *PR1* **(b)** were quantified using qPCR and were normalized to *Rubisco*. The expression levels are shown as $2^{-\Delta CT}$. The experiment was repeated three times. Error bars represent SEM. Statistical significance was determined using a Student *t*-test (unpaired two-tailed *t*-test, $P < 0.05$); ns, not significant.

**Table S1:** Nucleotide sequences of the primers that were used in this study.

| Primers for generating constructs | Sequence (5′ - 3′)† | Primer code |
|---|---|---|
| TOPO Cf-5:CO Fw* | caccATGGGATTTGTTCTCTTTTCACA | Eo_040 |
| TOPO Cf-5:CO Rv* | GAACCTATTGTTCCTCCGCC | Eo_041 |
| **Primers for qPCR** | | **References** |
| CfActin Fw | GGCACCAATCAACCCAAAG | |
| CfActin Rv | TACGACCAGAAGCGTACAG | |
| Rubisco Fw | GAACAGTTTCTCACTGTTGAC | |
| Rubisco Rv | CGTGAGAACCATAAGTCACC | Mesarich et al., 2014 |
| Avr4 Fw | CCCCAAAACTCAACCATACAAC | |
| Avr4 Rv | GCTTCGCATTGCCAACTTC | |
| Avr5 Fw | TGGTAGCTACGACGCCCTAC | |
| Avr5 Rv | TCGCATTGCTGATGGTCTAC | |
| SIPR1 Fw | TGGTGACTTCACGGGGAGGG | Zarate et al., 2007 |
| SIPR1 Rv | CGGACTGAGTTGCGCCAGAC | |

*Fw, forward; Rv, reverse.

†The sequence that is required for the TOPO reaction is shown in lowercase letters.
